# Thalamic deep brain stimulation as a paradigm to reduce consciousness: implications for cortico-striatal dynamics, absence epilepsy and consciousness studies

**DOI:** 10.1101/2021.07.27.453855

**Authors:** Michelle J. Redinbaugh, Mohsen Afrasiabi, Jessica M. Phillips, Niranjan A. Kambi, Sounak Mohanta, Yuri B. Saalmann

## Abstract

Anesthetic manipulations provide much-needed causal evidence for neural correlates of consciousness, but non-specific drug effects complicate their interpretation. Evidence suggests that thalamic deep brain stimulation (DBS) can either increase or decrease consciousness, depending on the stimulation target and parameters. The putative role of the central lateral thalamus (CL) in consciousness makes it an ideal DBS target to manipulate circuit-level mechanisms in cortico-striato-thalamic (CST) systems, thereby influencing consciousness and related processes. We used multi-microelectrode DBS targeted to CL in macaques while recording from frontal, parietal, and striatal regions. DBS induced episodes reminiscent of absence epilepsy, here termed absence-like activity (ALA), with decreased behavior and vacant staring coinciding with low-frequency oscillations. DBS modulated ALA likelihood in a frequency-specific manner. ALA events corresponded to decreases in measures of neural complexity (entropy) and integration (Φ*), an index of consciousness, and substantial changes to communication in CST circuits. During ALA, power spectral density and coherence at low frequencies increased across CST circuits, especially in thalamo-parietal and cortico-striatal pathways. Decreased consciousness and neural integration corresponded to shifts in cortico-striatal network configurations that dissociated parietal and subcortical structures. Overall, the features of ALA and implicated networks were similar to those of absence epilepsy. As this same multi-microelectrode DBS method – but at different stimulation frequencies – can also increase consciousness in anesthetized macaques, it can be used to flexibly address questions of consciousness with limited confounds, as well as inform clinical investigations of absence epilepsy and other consciousness disorders.

**SIGNIFICANCE:** We use tailored, multi-microelectrode thalamic deep brain stimulation to reversibly decrease consciousness for otherwise healthy, wakeful animals in a stimulation frequency-dependent manner. This represents a bidirectional mechanism for controlling consciousness, as the same method can increase consciousness under certain conditions. Theories of consciousness debate the relative contribution of parietal and frontal lobes, and largely ignore subcortical contributions. In this study, changes in consciousness predominantly involve changes in subcortical and parietal regions, implying that they contribute more to consciousness than frontal regions. Further, decreases in consciousness (indexed by Φ*) coincide with decreased movement, staring, and low-frequency activity in the EEG, similar to absence epilepsy. Thus, the systems-level mechanisms for decreased consciousness in this study have broader clinical implications for absence epilepsy.

## INTRODUCTION

Deep brain stimulation (DBS) is a powerful clinical tool to treat severe conditions resistant to other interventions, yet the mechanisms remain unclear (Chiken and Nambu, 2016). DBS may lead to neural excitation/inhibition, regularized activity or altered neural communication. Further, effectiveness varies depending on location, electrode type, and stimulation parameters (Perlmutter and Mink, 2006).

Intralaminar thalamic DBS can be an effective tool to increase consciousness in minimally conscious state patients (Schiff et al., 2007) and anesthetized primates (Redinbaugh et al., 2020; Bastos et al., 2021). Of the intralaminar nuclei, the central lateral nucleus (CL) appears the most promising target, with anatomical connectivity well-suited to modulate consciousness. Theories of consciousness implicate interactions within and/or between frontal and parietal cortex (Lamme, 2006; Friston, 2010; Dehaene and Changeux, 2011; Oizumi et al., 2014). CL projects to both, providing input to superficial layers, and reciprocally interacting with deep layers (Kaufman and Rosenquist, 1985; Towns et al., 1990), which have been shown to be critical for consciousness (Senzai et al., 2019; Redinbaugh et al., 2020; Suzuki and Larkum, 2020). CL also receives input from the reticular activating system and projects directly to the basal ganglia (Smith et al., 2004; Smith et al., 2014), serving as both a key input and output of the structure (Jones, 2007). While the role of the basal ganglia in consciousness is debated (Tononi, 2004; Schiff, 2010; Boly et al., 2017), the striatum contributes to integrated information (Afrasiabi et al., 2021), a measure of neural complexity associated with consciousness (Oizumi et al., 2014), and contributes strongly to decoding neural differences between conscious states (Afrasiabi et al., 2021). Further, the basal ganglia are linked to altered consciousness with hallucinogens (Preller et al., 2019) and are suppressed during general anesthesia (Mhuircheartaigh et al., 2010) and absence epilepsy (Berman et al., 2010; Carney et al., 2010).

Empirical evidence suggests CL manipulations can bidirectionally influence consciousness. CL lesions are linked to consciousness disorders, like spatial neglect and coma (Schiff, 2008), and thalamic DBS has been reported to induce sleep or produce a syndrome similar to absence epilepsy in cats (Jasper and Droogleever-Fortuyn, 1948), depending on the DBS method, e.g., stimulation frequency (Hunter and Jasper, 1949; Ingvar, 1955). In rodents, optogenetic stimulation of CL at 40Hz or 100Hz can drive arousal and activity in cortico-striatal-thalamic (CST) networks during sleep, while 10Hz leads to behavioral arrest (Liu et al., 2015). Similarly, using a multi-microelectrode stimulation method (rather than applying current via 1-2 large electrode contacts in conventional DBS) in macaques (Fig. 1), we have shown that DBS specific to CL, more than neighboring thalamic regions, awakens animals from anesthesia when applied at 50Hz, rather than 10Hz or 200Hz (Redinbaugh et al., 2020). These frequency-specific effects may relate to natural rhythms of CL. In cat (Glenn and Steriade, 1982; Steriade et al., 1993) and monkey (Redinbaugh et al., 2020), a subset of CL neurons exhibit high firing rate (around 50Hz) during wakefulness, but lower firing during sleep and anesthesia. Thus, frequency-specific effects with respect to consciousness may relate to stimulation parameters mimicking these firing patterns (Redinbaugh et al., 2020). We hypothesize that stimulation frequencies mimicking CL activity patterns during sleep/anesthesia, or inducing abnormal CL activity, will disrupt consciousness in awake primates.

**Figure 1.**
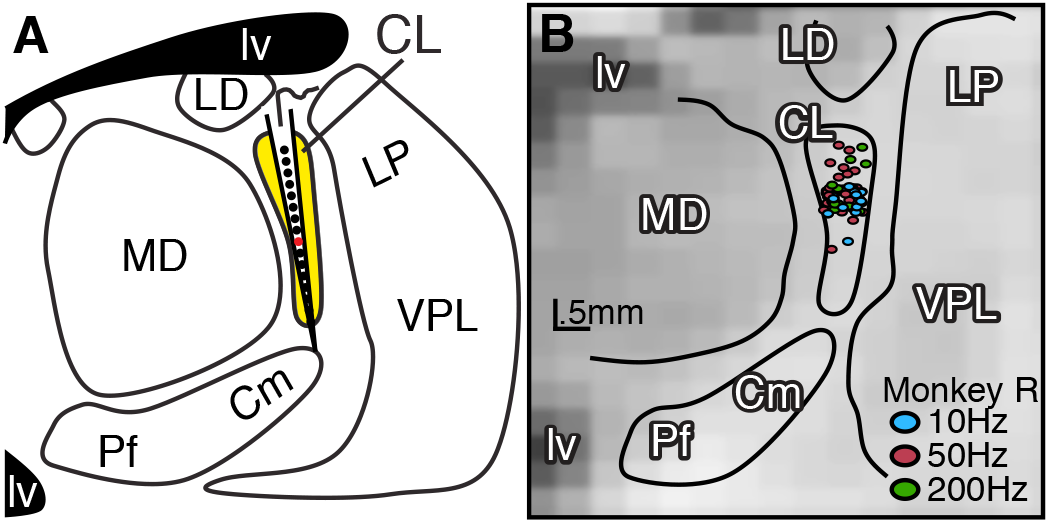
Structural imaging used to target multi-microelectrode DBS to the central lateral thalamus. ***A***, Schematic of coronal thalamic section (right hemisphere) with tailored DBS electrode placed such that contacts span the dorsal-ventral extent of CL (yellow). We simultaneously stimulated via 16 electrode contacts (200*μ*m spacing between contacts), with the centermost contact (contact 8) shown in red. ***B***, Track reconstruction of DBS sites overlain on the high-resolution structural image of the thalamus (monkey R, right hemisphere, A8). Only the centermost contact location shown (colored circles) for clarity. For each stimulation, 8 contacts above and 7 contacts below the colored circle would also be stimulated. To improve visualization by reducing overlap, positions plotted with up to 1 voxel (.5mm) jitter separately for 10, 50, and 200Hz experiments. Black lines demarcate different thalamic nuclei: central lateral, CL; centromedian, Cm; lateral dorsal, LD; lateral posterior, LP; mediodorsal, MD; parafascicular, Pf; ventral posterior lateral, VPL. Stimulation performed at all frequencies for most sites.

To test this hypothesis, we used a CL-tailored method of DBS in resting-state (wake) and rewarded (fixation task) conditions while simultaneously recording from frontal cortex, parietal cortex and striatum of macaques; we also recorded from CL in a DBS-free condition. We applied DBS at different frequencies (10, 50, 200Hz) across a series of stimulation experiments, as well as experimental series without DBS, to test effect specificity (Fig. 2). We expected that 50Hz DBS, mimicking wake-state CL activity patterns, would not affect waking behavior; but 10/200Hz DBS (atypical frequencies for wakefulness) would reduce consciousness. Moreover, reduced consciousness would relate to aberrant communication along CST pathways, thought to play a role in consciousness disorders (Schiff, 2010; Blumenfeld, 2012).

**Figure 2.**
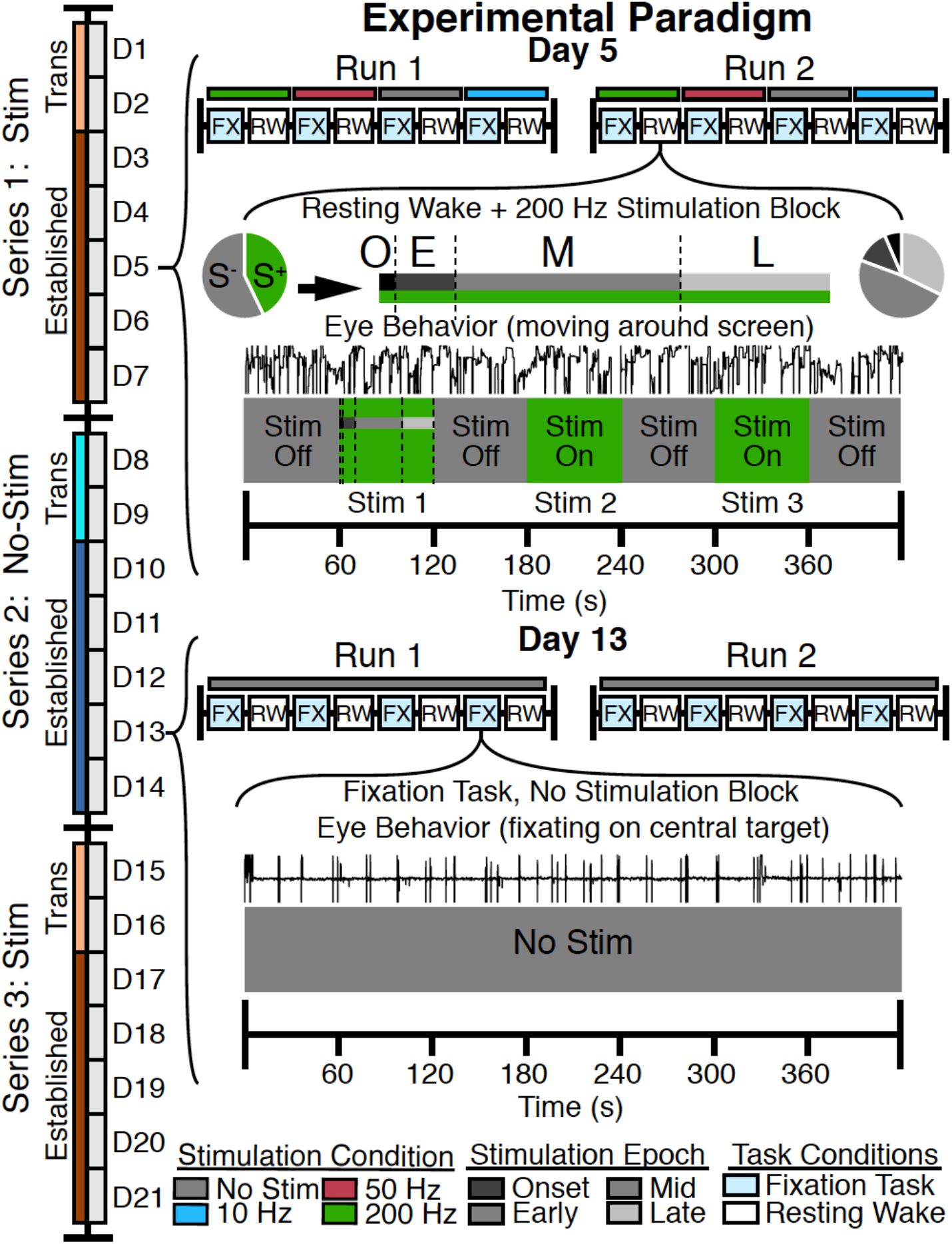
Experimental paradigm for manipulating consciousness in awake macaques. Schematized paradigm to reveal stimulation frequency-specific effects of thalamic DBS across multiple time scales. On the far left, a schematized timeline of the paradigm shown shifting between a series of experiments with (Stim) and without (No-Stim) DBS. The first two recording days of a series were considered transitional, and the following days in the same series were considered established. The top half of the figure presents an example day (D5) in the DBS paradigm, comprised of a pair of experimental runs with pseudorandom stimulation frequency assignment. Here, a sample resting-wake block (420s) using 200Hz DBS is featured, with typical on/off periods (S^+^/S^-^) of DBS within the block (60s on, 60s off). Above the block, one stimulation with duration of 60s is subdivided into shorter time periods: onset (O, 0-2s), early (E, 2-10s), mid (M, 10-40s), and late (L, 40-60s) periods with respect to the start of DBS. The bottom half of the figure presents an example day (D13) in the established No-Stim portion of the paradigm, with a sample fixation task block featured. Here, the eye-tracker trace indicates the animal spends most of the experiment fixating on the central target (centered, steady) as compared to the upper trace where the animal shows typical eye behavior during resting wakefulness (eyes moving around).

## MATERIALS AND METHODS

### Model

We acquired data from two male monkeys (*Macaca mulatta*, 4.3-5.5 years old, 7.63-10.30 kg body weight) housed at the Wisconsin National Primate Research Center (WNPRC). All procedures conformed to the National Institutes of Health Guide for the Care and Use of Laboratory Animals, and were approved by the University of Wisconsin-Madison Institutional Animal Care and Use Committee.

### Surgery and Electrode Placement

We performed a stereotaxic head implant and craniotomy surgery using aseptic techniques under isoflurane general anesthesia (1-2%) according to the procedures described previously (Redinbaugh et al., 2020; Afrasiabi et al., 2021). We placed 2.5mm craniotomies over the frontal eye fields (FEF), lateral intraparietal area (LIP), CL, and caudate nucleus (CN) using stereotaxic measurements based on high-quality structural MRI acquired in advance of the surgery and comparisons to a stereotaxic atlas (Saleem and Logothetis, 2007). We also acquired MRI subsequent to surgery with electrodes *in situ* to verify electrode placements. We used a GE MR750 3T scanner to acquire 3D T1-weighted structural images from the anesthetized monkeys, with an inversion-recovery prepared gradient echo sequence (FOV=128 mm^2^; matrix=256 × 256; no. of slices=166; 0.5 mm isotropic; TR=9.68 ms; TE=4.192 ms; flip angle=12°; inversion time (TI)=450 ms). We averaged 6-10 T1-weighted images for the pre-surgery high-quality structural image of each monkey, and we averaged 2 T1-weighted images to visualize electrodes *in situ*. Further imaging details have been described previously (Redinbaugh et al., 2020; Afrasiabi et al., 2021). Electrode placements were further verified online using functional criteria. Local electrical stimulation at low current in FEF produced eye movements (Bruce et al., 1985), many neurons in the LIP region of interest (ROI) exhibited the classic response characteristic of peri-saccadic activity, and the CL ROI had a subset of neurons with high firing rates around 40-50Hz as previously reported in CL (Glenn and Steriade, 1982). Finally, we performed post-mortem histology in one monkey, and visualized Hematoxylin and Eosin-stained sections (8 *μ*m) under a microscope, to verify electrode tracks in ROIs.

We separated cortical ROIs into superficial and deep layers using inverse current source density (iCSD) analysis of auditory-evoked ERPs (Pettersen et al., 2006). Both FEF (Schall et al., 1995; Romanski et al., 1999; Caruso et al., 2016) and LIP (Cohen et al., 2005; Gifford and Cohen, 2005) have been shown to respond to auditory stimuli. The demarcation between superficial and deep layers was the bottom of the earliest sink after auditory stimulation. We then assigned electrode contacts to iCSD-defined superficial and deep layers, and confirmed layer assignments using reconstructions of recording sites along each electrode track (computed from electrode depth measurements and MRI of electrodes *in situ*) and spiking activity (demarcating gray from white matter). We used auditory-evoked ERPs to allow comparison of wake data with anesthesia data. CSDs were computed using the CSDplotter toolbox (https://github.com/espenhgn/CSDplotter) for MATLAB (dt = 1 ms, cortical conductivity value = 0.4 S/m, diameter = 0.5 mm). Tones were played in a sequence (200 ms duration; 800 ± 100 ms jitter between tones) comprised of 80% standard tones (0.9 kHz) and 20% deviant/oddball (1 kHz). Further details have been previously described (Redinbaugh et al., 2020; Afrasiabi et al., 2021).

### Electrophysiological Recordings and Deep Brain Stimulation

We recorded broadband neural activity (filtered 0.1-7,500Hz, amplified and sampled at 40kHz) using a preamplifier with a high input impedance headstage and OmniPlex data acquisition system controlled by PlexControl software. Probes were MRI-compatible linear microelectrode arrays (MicroProbes) with 16 (FEF and CN) or 24 (LIP and CL) contacts; all were platinum/iridium electrodes 12.5*μ*m in diameter with 200*μ*m spacing between contacts and impedance between 0.8 and 1.0 MΩ. We recorded EEG with titanium skull screws located above dorsal fronto-parietal cortex.

We performed DBS using electrode arrays (24-contacts, 200*μ*m spacing) previously used as recording electrodes, which have reduced impedance. Stimulations (400*μ*s biphasic pulses, negative phase first) occurred simultaneously across the deepest 16 electrode contacts for 60s at 10, 50 or 200Hz with 200*μ*A of current. These parameters had been previously titrated and successfully used to influence consciousness (Redinbaugh et al., 2020).

### Experimental Paradigm

There were 40 total 2-4 hour recording days (18 for monkey R; 22 for monkey W). Monkeys sat upright in a primate chair in a quiet dark room. The animal’s head was immobilized using a head post and/or four rods that slid into fabricated hollow slots in the acrylic implant. Each day, we inserted electrodes into ROIs (CL, CN, FEF, and LIP) through chronically implanted guide tubes (Redinbaugh et al., 2020).

We exposed animals to a paradigm that included our thalamic DBS method, which involved a series of recording days in which animals received DBS (Stim) and separate series of recording days in which animals did not receive DBS (No-Stim). The first two days after switching (as well as weekends, when recordings did not take place) between these series types were considered transitional, and all other days in the same series were considered established, as this allowed sufficient data for analysis. Figure 2 depicts the schematized paradigm based on Monkey W. The paradigm was similar for monkey R, but started in the No-Stim condition following previously described anesthetized recordings and DBS experiments (Redinbaugh et al., 2020). During a non-stimulation series, recordings included 4 brain areas (CL, CN, FEF, LIP). During a stimulation series, we applied DBS through the CL probe, precluding acquisition of CL recordings. On each recording day, monkeys alternated between task-on (rewarded fixation task, “FX”) and task-off (resting wake, “RW”) blocks. In stimulation series, each pair of task on/off blocks included thalamic DBS pseudorandomly assigned to one of four conditions (10/50/200 Hz stimulation frequency and No-Stim control). After completing a run of all conditions (8 blocks, i.e., task on/off at 10Hz, 50Hz, 200Hz and No-Stim control), we started another run. Each block of about seven minutes duration included three stimulation replications: typically, stimulations occurred 1-2, 3-4 and 5-6 minutes after the start of the block, with one minute allowed inbetween for recovery. To characterize the timing of observed effects during the 60s stimulations, we considered specific time windows relative to the onset of DBS: onset (within 2s of the start of DBS), early (2-10s since start), mid (10-40s since start), and late (40-60s since start). Onset thus described the region of stimulation very close to initiation. In its entirety, this paradigm allowed us to investigate effects at the level of individual stimulations, blocks, recording days, and across the entire history of Stim and No-Stim series including transitions between them.

For the general anesthesia data in Figures 5 and 6, we used either isoflurane (0.8-1.5% on 1 L/min O_2_ flow; 9 sessions: 5 for Monkey R, 4 for Monkey W) or propofol (0.17-0.33 mg/kg/min i.v.; 9 sessions: 4 for Monkey R, 5 for Monkey W). Full details have been previously described (Redinbaugh et al., 2020).

### Behavioral Monitoring and Video Processing

During recordings, experimenters closely observed animal behavior. We monitored eye movements using an eye-tracker and videos focused on the animal’s face recorded evidence of facial (mouth and nose) movements and eye openness. Offline, videos were processed using MATLAB and converted into continuous luminance traces based on the average luminance in small windows around the animal’s eyes, mouth, and nose. Eye traces revealed periods of open (low luminance) and closed (high luminance) eyes. Derivatives of luminance around the mouth and nose revealed periods when the mouth and nose were moving, or when movement was reduced.

### Absence-Like-Activity Detection

DBS often triggered long-duration periods of behavioral arrest very similar to absence epilepsy, and thus we refer to these events as absence-like activity (ALA) in this study. Absence epilepsy is hallmarked by brief (10s on average) and sudden loss of consciousness, without convulsive movements. Humans often stare vacantly during an episode and motor movements are either reduced (typically without falling down) or very small and repetitive in nature (automatisms, for example, stereotyped small lip smacks) (Blumenfeld, 2005; Hughes, 2009; Tenney and Glauser, 2013).

For systematic detection of ALA, we used custom code identifying periods of time when eye movement was decreased (based on the derivative of the polar coordinates of the eye tracker data), eyes remained open and not partially lidded (based on video eye traces), and facial movement was reduced and/or small and repetitive (based on video mouth and nose traces) (see Fig. 4*A,B*). Onset times were defined based on when the changes in behavior occurred (stillness and little/no eye movement) and offset was defined by when normal behavior resumed (eye/face/limb movement resumed). Due to technical issues, eye tracker data were not recorded on a subset of ALA events (n=4), and video data were not recorded on a separate subset (n=3). Detection was then based on the remaining signals, as well as behavioral notes. Due to the high degree of correlation between these metrics (Fig. 4A), this did not substantially affect detection. We defined pre-and post-event conditions as the entire time series up to 30s before or after event onset or offset, respectively. We identified possible events as those lasting for at least 4.2s, which exceeded the 95% confidence interval of fixation durations based on the rest of the awake data set, and if the location of the animal’s gaze did not match the location of the central fixation target in the fixation task (within 3° of visual angle).

Absence is associated with low-frequency oscillatory activity between 2 to 10 Hz depending on the species (Coenen and Van Luijtelaar, 2003; Blumenfeld, 2005; Hughes, 2009; Tenney and Glauser, 2013). Online, many of the observed events coincided with low-frequency EEG activity. Offline, we assigned events as ALA if they involved significant increases in low-frequency activity at any frequencies 1-9Hz in the EEG relative to the pre-ALA condition. A range of frequencies was used because the oscillatory frequency associated with absence epilepsy has been known to start higher and reduce over the time course of events (Steriade, 1974; Bosnyakova et al., 2006; Bai et al., 2010). Further, we wanted to ensure that our analyses were not biased towards any one possible frequency common to the absence range. In contrast, a subset of the automatically detected stable eye periods showed no evidence of atypical low-frequency activity. These were considered to be natural stares and used as the primary control in this study (see below for additional controls), as they shared similarities in behavior to ALA.

### Controls

To identify neural correlates of ALA, we compared neural activity during ALA to a variety of controls accounting for activity differences which could be attributed to behavior, consciousness, or the typical dynamics of wakefulness and attention. To account for time-specific differences, we compared ALA to data recorded 30 seconds prior to (Pre) or after (Post) event onset/offset. To account for differences related to consciousness but not behavior per se, we compared ALA to resting wake data, when the animals were conscious but not performing a specific task. Resting data derived from epochs when eye position was stable (for at least 1s), and eyes were open (other face or body movements may have occurred during these windows). The resting state controls were also case-matched (data taken from same recording day with same task, DBS type, and electrode positioning) to account for the experimental paradigm (Stim/No-Stim and stimulation frequency) and electrode positions. ALA was associated with staring. To account for possible goal-directed fixations, ALA was compared to data from a fixation task, where animals received juice rewards the longer they fixated on a faint, central target. Facial movements were common as animals licked juice rewards. Again, controls were case-matched and we only analyzed data during which the eyes were open and the animal was fixating. The fixation task also allowed comparisons between neural mechanisms present during resting wakefulness and cognitive engagement, which both reflect consciousness. Finally, to account for differences in consciousness while controlling for behavior, we compared ALA to long-duration stares without atypical low-frequency EEG activity (stare, STR). While these control stares could not be case-matched, as they were rare and occurred randomly in the data set, they could be temporally aligned, similarly to ALA, with a defined onset, offset, pre and post period. This allowed us to compare the temporal dynamics of ALA directly to highly-specific controls.

### Local field potential (LFP) and spike preprocessing

For ALA and stare events, we extracted data up to 30 seconds before behavioral onset (Pre), and up to 30 seconds after offset (Post) when possible. For resting wake controls and the fixation task, we first determined stable eye epochs, to match eye behavior between different conscious states (wakefulness, ALA, general anesthesia). These epochs started 200ms after a saccade and ended 200ms before the next saccade. For all events and controls, we divided epochs of data into non-overlapping 1s windows for analysis, which can be considered similar to trials. For LFPs, we bandpass filtered data from 1-250Hz (Butterworth, order 6, zero-phase filter) and downsampled to 1000Hz sampling frequency. We linearly detrended LFPs, then removed powerline noise (significant sine waves at 60Hz) using the function rmlinesc from the Chronux data analysis toolbox for MATLAB (http://chonux.org/). When individual electrode contacts had signal amplitude greater than 5 standard deviations from the mean of other contacts on the same probe, they were excluded from the analysis. For power spectral density and coherence analyses, we calculated bipolar derivations of the LFPs (the difference between adjacent electrode contacts) to curtail possible effects of a common reference and volume conduction (Bollimunta et al., 2008; Haegens et al., 2015; Trongnetrpunya et al., 2015). Prior to entropy (*H)* and Φ* analyses, we performed ICA (Independent Component Analysis) to minimize the linear dependencies among LFP contacts in each area, then we normalized each LFP to its mean across all epochs for that recording day. We then binarized each LFP with respect to its median amplitude over the 1s epoch, to remove potential biases related to amplitude differences across channels or conditions.

For spiking activity, we bandpass filtered data from 250-5,000 Hz (Butterworth, order 4, zero-phase filter) and sorted spikes using Plexon Offline Sorter software. Spikes were initially detected with thresholds greater than 3 standard deviations from the mean. We used principal components analysis to extract spike shape features and the t-distribution expectation maximization estimation algorithm to determine clusters of spikes with similar features.

During DBS, stimulations induced a brief artifact caused by the applied current. To remove this artifact, we excised a 1ms window around the artifact, linearly interpolated across this window, and removed any significant sine waves at the stimulation frequency using the Chronux function rmlinesc.

### Spike rate

We calculated the average spike count across the same 1s windows as the LFP analyses. For epochs that included thalamic DBS, artifact removal decreased the viable window as a function of the stimulation frequency. Thus, the window size was adjusted accordingly and accounted for in the spike rate calculation. For each neuron, spike rate was normalized to data collected during the pre-event condition, to allow comparison with all possible controls. We only analyzed neurons if they were recorded during an ALA or control stare event (in addition to other controls).

### Power and coherence spectrograms

For ALA and control stare events, we calculated power spectrograms using 1s sliding windows with 0.1s stride/overlap across the entire time course of the event aligned to the onset of the behavior. Spectrograms were calculated for every bipolar-derived LFP using multi-taper methods (5 Slepian taper functions, time bandwidth product of 3) with the Chronux function mtspecgramc and then averaged across all viable electrode contacts.

To measure the temporal relationship between LFPs, we used coherence within and between CL, CN, FEF and LIP. Coherence is given by:

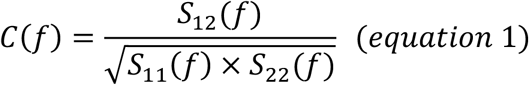

where S(f) is the estimated power spectral density with subscripts 1 and 2 referring to the simultaneously recorded LFPs at two different sites. We applied multi-taper methods (5 Slepian taper functions, time bandwidth product of 3) to yield coherence spectrograms, using the Chronux function cohgramc. For ALA and control stare events, we calculated coherence in 1s sliding windows, stepped 0.1s, for all bipolar-derived LFP pairs corresponding to anatomically plausible pathways. We then averaged coherence values for all LFP pairs representing each pathway.

#### Φ* Computation

F is a proposed index of consciousness measuring both the complexity and integrated structure of a given system (Oizumi et al., 2014; Oizumi et al., 2016). We calculated Φ* (Barrett and Seth, 2011; Oizumi et al., 2016; Haun et al., 2017; Afrasiabi et al., 2021) as a practicable estimate of integrated information (F), because directly calculating the Φ of large systems is intractable (Oizumi et al., 2016). To compute Φ* based on LFPs, we formed the state of a subsystem *X*(*t*) at time *t* (1 ms time bins considering 1 kHz sampling frequency) as:

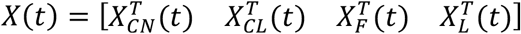

where its elements are the bipolar-derived LFP signals, ranging from 1 to 23 signals (resulting from 24 electrode contacts) in each brain area: CN, CL, FEF (F) and LIP (L). The component of *X*(*t*) for each area is *N*_*ch*_ × *T*-sized, where *N*_*ch*_ *s*pecifies the number of bipolar-derived channels for each of four areas and *T* = 1000 (1s windows, sampled at 1kHz). We next calculated the uncertainty about the states assuming a multivariate Gaussian distribution of the states across time as:

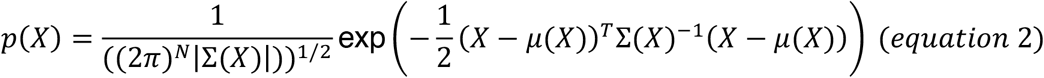

where Σ(*X*) (*t* removed to be concise) is the covariance matrix of 5 estimated over a 1s data window, *μ*(*X*) is the mean of the state vector 5 over the window, |Σ(*X*)| is the determinant of the covariance matrix that can be considered a measure of uncertainty about the state *X* at any time point within the window, and *N* is the total number of channels across areas. We used the Henze-Zirkler Multivariate Normality test (Henze and Zirkler, 1990) to verify that our states had multivariate Gaussian distributions.

The entropy for the states *X*(*t*), given its probability density function (pdf) as *p*(*X*), is defined as:

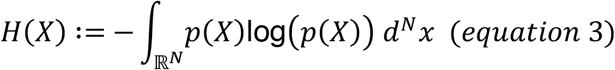

The entropy is maximized if *p*(*X*) *is* multivariate Gaussian, and can be calculated in closed form as:

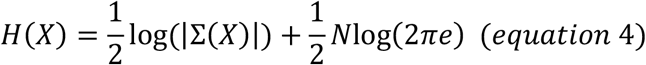

which can be described as the uncertainty about state *X*(*t*) at time *t*. The mutual information, i.e., the reduction of uncertainty about state *X*(*t*) at time *t*, given its past at time *t − τ* can be calculated as:

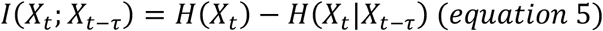

Where *H*(*X*_*t*_|*X*_*t*−*τ*_) is the conditional entropy of state *X*(*t*) given its past *X*(*t − τ*), and can be derived in canonical form with a Gaussian distribution assumption of states as:

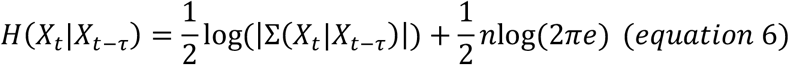

where Σ(*X*_*t*_|*X*_*t*−*τ*_) is the covariance matrix of the conditional distribution *p*(*X*_*t*_|*X*_*t*−*τ*_) (conditional covariance) that can be expressed analytically as:

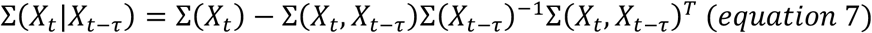

where Σ(*X*_*t*_|*X*_*t*−*τ*_) is the cross-covariance matrix. Mutual information, *I*, can be regarded as a measure of the information the current state has about its past, and it is used to calculate Φ*, a measure of integrated information. Φ* of the subsystem *X*(*t*) is information that cannot be partitioned into independent parts of *X* and can be defined as:

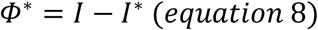

Where *I*\* (disconnected *I*) is called mismatched information. We calculated *I*\* for every bipartition of system *X*. System parts of interest were defined CN, F_s_, F_m_, F_d_, L_s_, L_m_, L_d_; where s, m and d subscripts correspond to superficial, middle and deep layers respectively. Within each cortical layer (F_s_, F_d_, etc.), there were multiple bipolar-derived channels. We did not partition these individual layers, and so each layer was effectively a subsystem part consisting of multiple parallel channels.

The partition *p* that minimizes the normalized Φ* is the *minimum information partition* (*MIP*), as defined in (Balduzzi and Tononi, 2008):

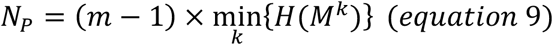

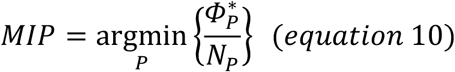

Here *m* is the number of partitions and *m*^*k*^ is the *k*^*th*^ part of subsystem *X. N*_*p*_ counterbalances inevitable asymmetries introduced by computing Φ* across a number of partitions of unequal sizes. The MIP is the weakest link between the parts of *X*, where its partition into subsystems results in minimal information loss. We calculated the covariance and cross-covariance matrices for each 1s window using the shrinkage method for a more stable result and averaged them across all windows for each recording day, to calculate the *MIP* and its corresponding Φ*, using gradient ascent and L-BFGS optimization method. We incorporated 2 NVIDIA GTX 1080ti GPUs to accelerate the process of searching for the MIP to calculate Φ*. We calculated *I* and Φ* with timelag τ = 15 *ms*, as this lag maximized Φ* in our previous study of this same CST system (Afrasiabi et al., 2021).

Φ* was computed using all available epochs for each condition (pre, post, ALA, control stare, resting wake, and fixation). Because Φ* is a relative metric that cannot be easily compared between systems with different composition (i.e., different brain areas), we excluded recordings where a given brain region was absent (for example, when no electrode contacts were identified in the deep cortical layers). To account for small differences in Φ* related to electrode placement, Φ* was centered relative to the values obtained during recording tasks in the paired fixation experiments. We also computed Φ* across time for ALA and control stare conditions, aligned to the onset of the event. In this case, Φ* was normalized (Z score) across all available time and then averaged for ALA and control stare events. For both centered and Z-score values, higher values of Φ* indicate increased consciousness, while lower values indicate decreased consciousness.

### Machine Learning

We computed the power spectrogram of bipolar derived LFPs in 1s epochs (0.5s sliding window) from 11 seconds prior to ALA or control stare onset up to 12 seconds after offset (5 Slepian tapers). Data were only further analyzed between -10.5 to 4.5 seconds (relative to event onset), as this represented the minimum consistent aligned segments across all events. Power values were averaged for each frequency band (delta 1-4 Hz; theta, 4-8 Hz, alpha, 8-15 Hz; beta, 15-30Hz; gamma 30-90Hz) and all contacts in each area of interest (CN, FEFs, FEFd, LIPs, LIPd) which resulted in 25 input features for each 1s time bin.

We trained a Bayesian classifier (Bishop, 2006) for each 1s time bin to classify the data into two states: ALA and Control Stare. For the Bayesian classifier, the class-conditional densities were modeled as gaussian distributions with equal covariances. We used an uninformative prior for classes (Normal-Jeffreys prior (Murphy, 2012)) to avoid any bias toward any of the classes. Given *X*_*t*_ as input feature vector at time *t*, the posterior probability that the data belongs to class c can be derived as follows:

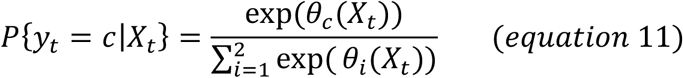

Where θ_*c*_(*X*_*t*_) will be calculated at each time bin 0 as:

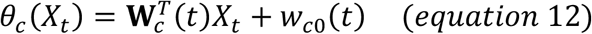

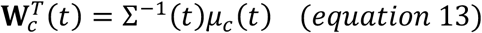

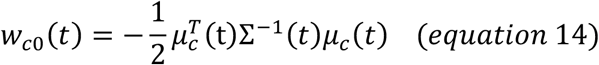

Where *μ*_*c*_(*t*) is the expected value of class *c* at time *t*; and ∑(*t*) is the common covariance matrix for all classes, which is calculated as the unweighted mean of two covariance matrices for two classes and subsequently regularized using shrinkage where the regularization parameter was derived during training and cross-validation. All the parameters in equations above were calculated using the training set at each fold of cross-validation.

We used 10-fold cross validation with 60% of the data as training set and 40% as test set. To test feature importance, we computed Mean Decrease in Accuracy (MDA). One at a time for each feature, we randomly permuted the sample labels (1000 times) and compared average accuracy to that of the unpermuted model. Permutation effectively erases the relationship between the given feature, condition, and other features, and thus the drop in accuracy can be interpreted as the relative importance of the permuted feature (Molnar, 2019).

### ALA Probability and Causal Power

To compare the relative effects of different stimulation frequencies, we computed ALA and control stare probability by recording block for each condition. To compare DBS effects across longer time scales (different experimental days) we computed ALA and control stare probability in sliding windows (step size = 1 block) consisting of 32 recording blocks (the typical number of blocks recorded in two days). Stimulation predominance (the number of blocks that included thalamic DBS over the 32 recording blocks) was also calculated in each window. Both metrics were normalized to the maximum for correlation analysis.

We computed causal power (CP) (Griffiths and Tenenbaum, 2005), for probabilities that represented a theoretical causal structure between a dependent outcome (ALA event) and an independent experimental variable (stimulation frequency). CP is a differential probability ratio indicating the degree to which a proposed cause predicts an outcome more so than alternative causes, and is calculated based on the conditional probability matrix of event (e) co-occurring with/out cause (c), as seen in equation 15. Positive/negative CP indicates that the probabilistic relationship is causal, and the proposed cause is more/less likely to lead to an event than alternatives.

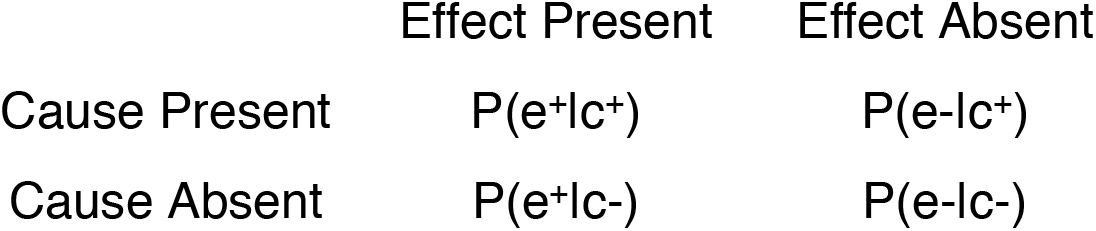

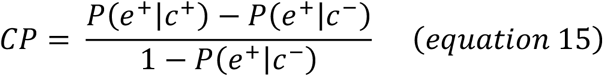

### Statistical Analyses

We performed statistical analyses using correlations and linear mixed effect models in R, the PyMC3 (Salvatier et al., 2016) package in Python, and MATLAB. We controlled all p-values (p) using the Holm-Bonferroni method, and report them as p_c_ when applicable. Models are shown for all interaction tests.

ALA probabilities (with respect to stimulation onset and frequency, Fig. 7*A,B,F*), causal power (Fig. 7*E*), and MIP association probabilities (Fig. 6*C,D*) were compared to their generated null-distribution (permutation test, see Fig. 3 for an example). We generated distributions (Fig. 3*A*) to represent a given population in our study, containing N elements with different member components (for example, N_A_, N_B_, N_C_ and N_D_ could represent the total number of experiments with each stimulation condition (10/50/200Hz and NS control), and n observations (n_A_, n_B_, n_C_ and n_D_ could represent the total number of ALA events observed with each condition). From that distribution, we drew n simulated random samples (Fig. 3*B*), without replacement if the population was finite and unrepeatable, or with replacement if otherwise, from the population, where n = the total number of observations. The process was repeated Nboot times, yielding a distribution of plausible observations under the null hypothesis of the given metric for each member category. Bootstrapping was always performed with replacement. Actual observations were then tested against these distributions to yield Z statistics and p-values (Fig. 3*C*), which were corrected for multiple comparisons. Comparisons of probability (Fig. 3*D*) and sample proportion (Fig. 3*E*), for example, can then be represented across member conditions relative to the null hypothesis, with error bars derived from the standard deviation of null distributions. Note, with this approach, increasing Nboot can increase the goodness of fit between the discrete distributions (Fig. 3*C*, bar histograms) and the fitted distribution (Fig. 3*C*, colored curves). However, the shape of the distribution, and thus the estimated error depends on the dimensions of the source population (N) and the sample rate (n). Because those values match what was done in the study, the resulting error bars reflect a reasonable estimate of the true, expected variance under the null hypothesis and cannot be inflated by Nboot.

**Figure 3.**
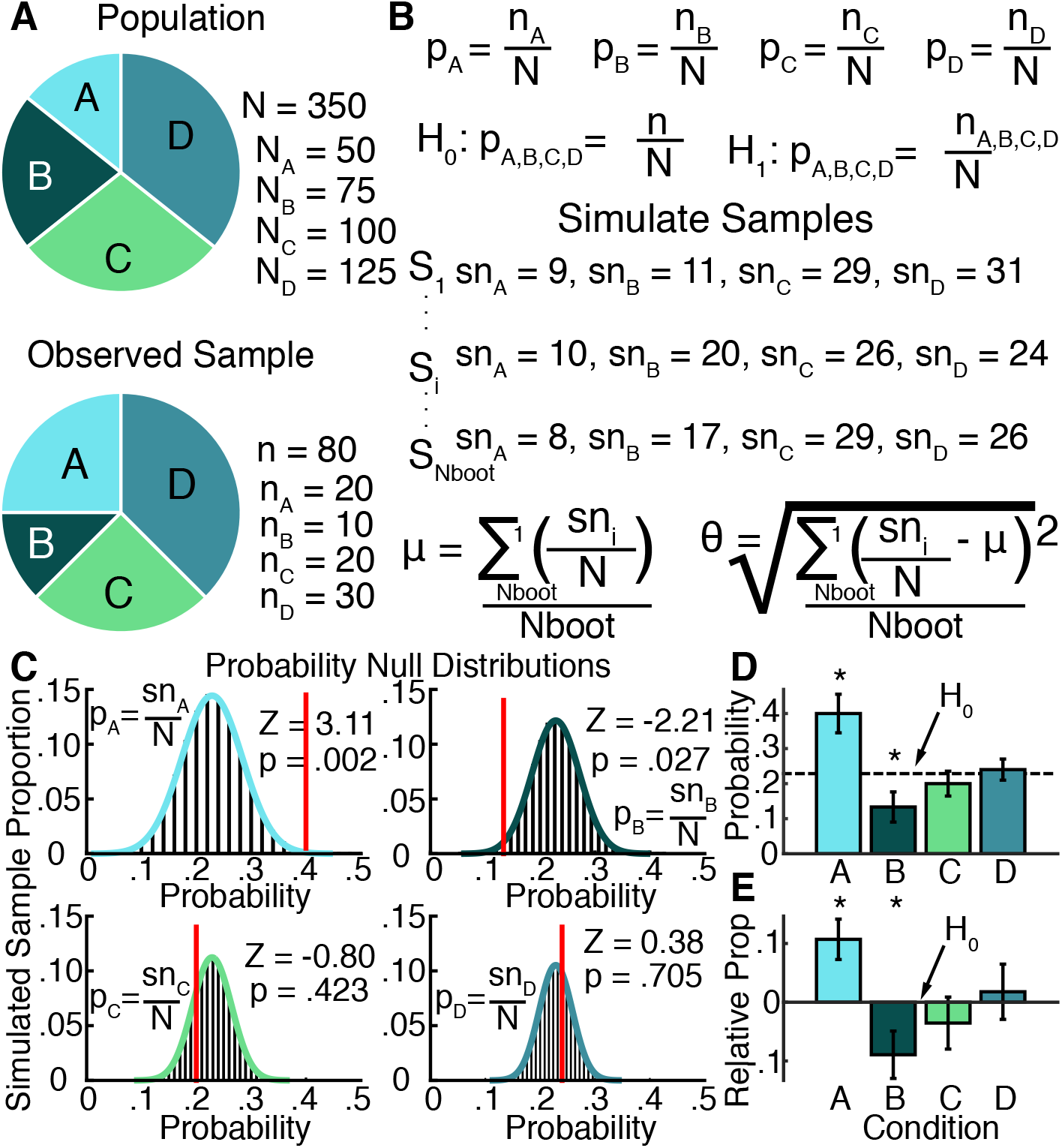
Example of simulated null distribution permutation test for probability analysis. ***A***, Example of hypothetical population, size N, consisting of 4 members and yielding observed sample, size n, consisting of the 4 members. The question to address: Is there category-specific variation in the probability of occurrence for different members of the observed samples? ***B***, Equations demonstrating how to represent the question in A as a null hypothesis, and how to simulate samples from the population. ***C***, Distributions (histograms) and fitted normal distributions (curves) showing the proportion of simulations yielding a given probability of member occurrence under the null hypothesis. Red lines indicate where the observed sample falls relative to the null distribution, with matching Z statistic and p-value before correction. ***D***, Category-specific probability of occurrence where error bars indicate ± 1SD of the null distribution. * indicates statistical significance. ***E***, Similar figure to (**D**), but representing the relative proportion of sample occurrences and the null hypothesis that the member proportions observed are the same as those observed in the population.

**Figure 4.**
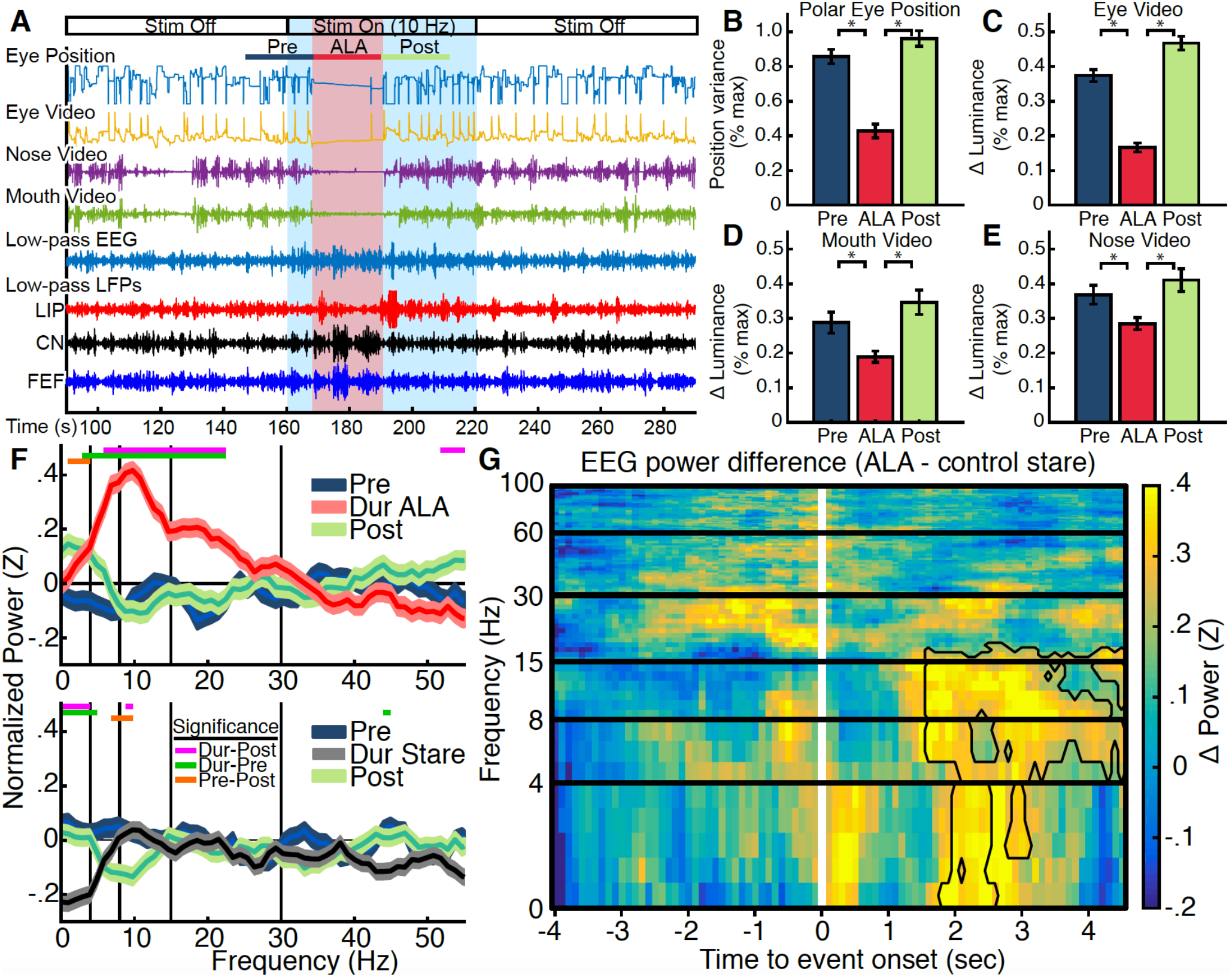
Thalamic DBS triggers periods of behavioral stillness coinciding with low-frequency activity in the EEG, similar to absence epilepsy. ***A***, Example time course of behavioral and neural signatures of an ALA event initiating after 10Hz stimulation. **B**,**C**,**D**,**E**, Average behavioral signatures during ALA relative to pre and post conditions (±SE) for the (**B)** variance in polar eye position from eye tracker, and change in luminance for video data centered on the (**C**) eyes, (**D**) mouth, and (**E**) nose. Luminance changes indicate movement. ***F***, Average EEG power (±SE) for ALA and control stares compared to pre and post conditions. Colored straight lines at top indicate regions of significance for t-tests across frequencies comparing spectra from the different conditions. Vertical black lines indicate the cutoff mark of different frequency bands: d = (0,4),θ = (4,8), α = (8, 15), β = (15, 30), *μ*_L_ = (30, 60), *μ*_H_ = (60, 100). ***G***, Spectrogram of average differences in normalized EEG power (ALA – control stare) aligned to event onset (white line). Frequency is represented across described power bands. Significant cluster outlined in black.

**Figure 5.**
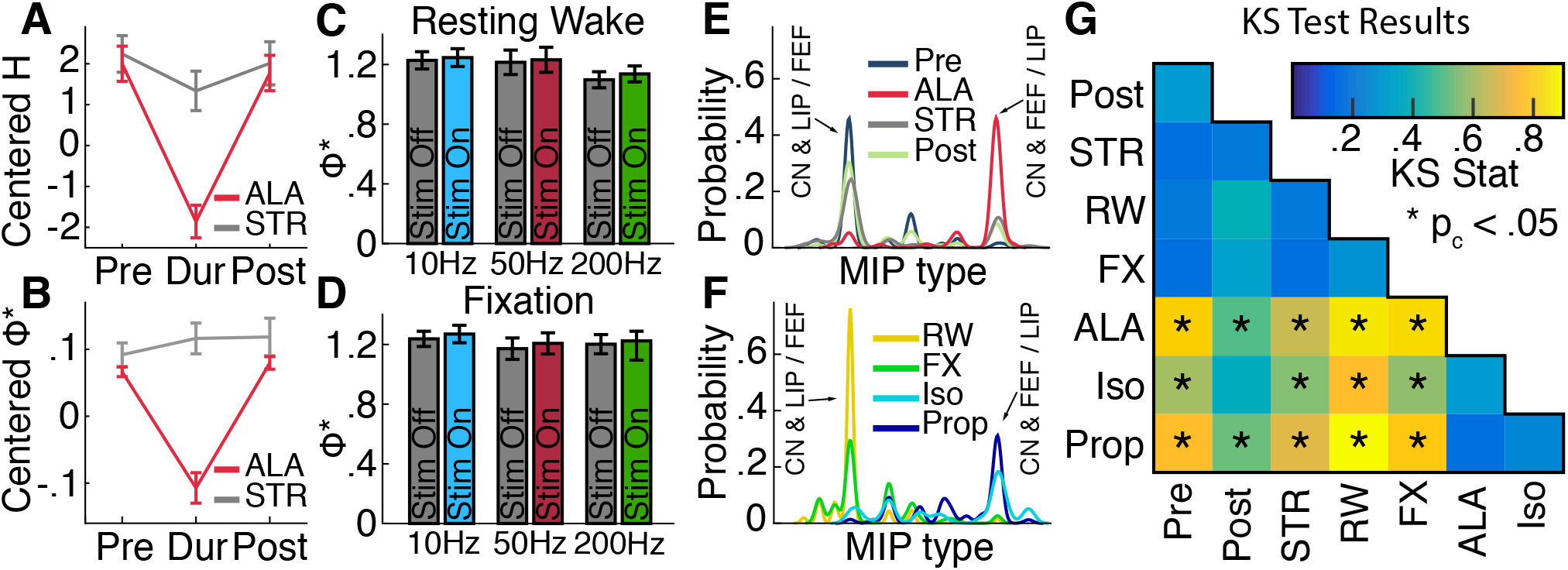
Measure of neural complexity and integration (Φ*) associated with consciousness selectively decrease during ALA, and integration patterns shift to reflect anesthetized rather than conscious states. ***A***, Average entropy (*H*) and ***B***, Φ* (±SE) centered relative to the fixation task for ALA and control stares relative to pre and post event conditions. ***C, D***, Average Φ* (±SE) for stim-off (gray bars) and stim-on (colored bars) epochs at different frequencies excluding ALA for resting wake (**C**) and fixation (**D**) controls. ***E, F***, Gaussian kernel-density estimates fitted to the probability distribution for the occurrence of different MIP types of the full system for (**E**) pre, ALA, control stare (STR), post, (**F**) resting wake (RW), fixation (FX), isoflurane (Iso), and propofol (Prop) conditions. ***G***, Results for the KS tests comparing the MIP kernel-density estimates across all conditions. Color scales with the strength of the KS stat. * indicates corrected p-value (p_c_) < .05.

**Figure 6.**
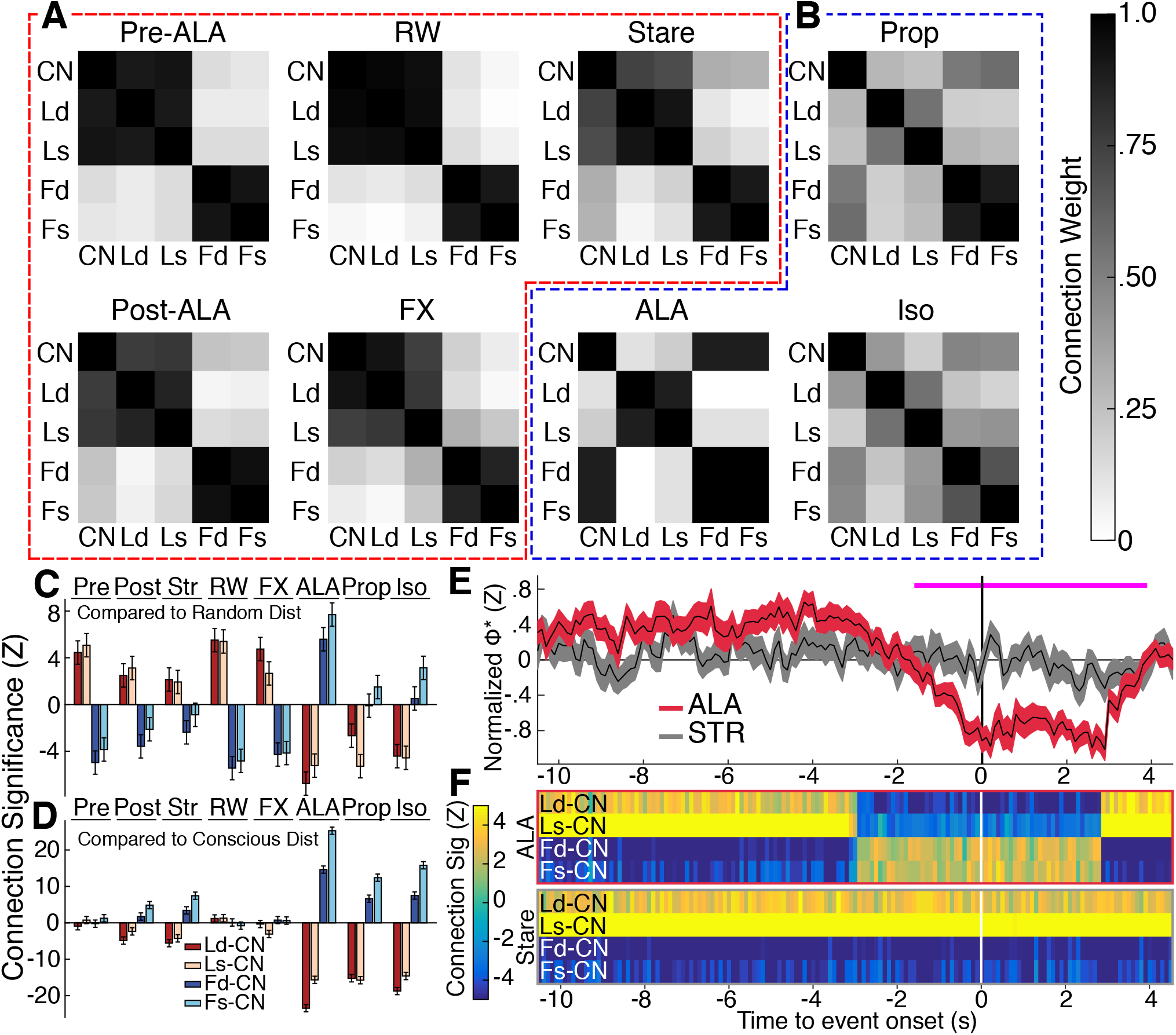
MIP changes reflect switches in parieto-striatal association indicative of consciousness. ***A, B***, Probability (connection weight) of each brain area associating with any other on the same side of the MIP for states assumed to be (**A**) conscious (pre-ALA, post-ALA, resting wake (RW), fixation task (FX) and control stares) or (**B**) less conscious (ALA, propofol (Prop), or isoflurane (Iso)). ***C, D***, Results (Z statistics ± SD of the null distribution) of the permutation test comparing cortico-striatal MIP associations under the null hypotheses that (**C**) MIP types occur randomly, or (**D**) MIP types reflect the same patterns as conscious states (defined by the resting wake and fixation task samples). Both approaches separate more consciousness from less conscious states. ***E***, Normalized Φ* (±SE) for ALA and control stares (STR) calculated in 1s sliding windows (.1s steps) and aligned to event onset (black vertical line). Horizontal pink line shows regions of significance for the pairwise t-test across time comparing ALA Φ* at each sample to the maximum. No significant differences were found in the control stare condition. ***F***, Cortico-striatal significance (Z) based on the sliding window approach in E for ALA and stare conditions. Computations used the same approach as in C, but now for each sample in the sliding window.

**Figure 7.**
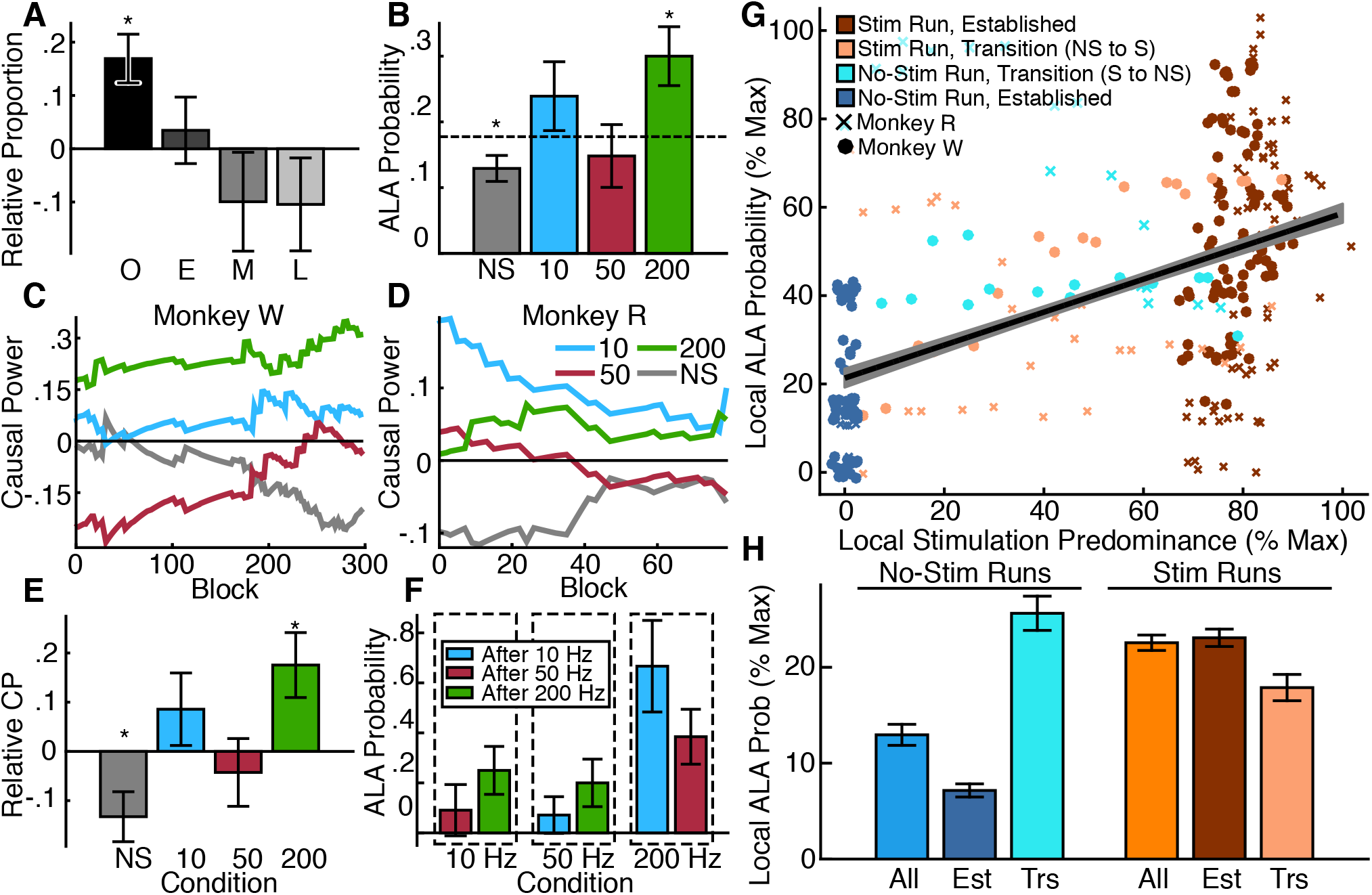
Thalamic DBS modulates ALA occurrence over acute and long time frames. ***A***, Relative proportion (± SD of the null distribution) of ALA events occurring around the onset (O, within 2 secs of stimulation start), early (E, 2-10s), middle (10-40s), or late (40-60s) phase of the 60s stimulations. ***B***, Probability of ALA (± SD of the null distribution) across different experimental conditions: no stimulation (NS) and stimulations at 10, 50 and 200Hz. Dashed line indicates the average probability of ALA (the null hypothesis). ***C, D***, Causal power by stimulation condition across experimental history (x axis starting when there are at least 10 blocks per condition to ensure robust causal power estimates) for (**C**) Monkey W and (**D**) Monkey R. ***E***, Population CP (± SD of the null distribution) under the null hypothesis that CP is equal across conditions and no different from 0. ***F***, Conditional probability (± SD of the null distribution) of ALA occurring with a particular stimulation frequency (10, 50 or 200Hz), given it was preceded by a different stimulation condition, under the null hypothesis that probability of ALA is consistent irrespective of preceding stimulations. ***G***, Correlation (best fit line ± SE of the point estimate) of local ALA probability and stimulation predominance (% maximum, sliding window across 32 recording blocks). Individual data points shown for both animals (R and W) across stimulation and no-stimulation runs during transitional (first two days) and established (third day onwards) phases of an experimental series. ***H***, Local ALA probability (± SE) for all data (All) in stimulation or no-stimulation runs, and further separated into established (Est) and transitional (Trs) phases.

To compare *MIPs* across states, we used a kernel density approach to estimate the probability mass functions for each MIP distribution for each state. We then applied a two-sample Kolmogorov-Smirnov test using the Pymc3 package in Python to determine whether these probability mass functions were different. We also compared the probability of different brain areas associating on the same side of the MIP for each state. These probabilities were then compared against the null distribution with 10,000 bootstraps.

We compared EEG power, LFP power, and LFP coherence time-frequency spectrograms between ALA and control stare conditions using the permutest function in MATLAB (Gerber, 2021), based on the Maris-Oostenveld method (Maris and Oostenveld, 2007). Tsum statistics, electrode counts, p-values, and bootstrap parameters are reported for significant clusters. We also performed t-tests with multiple comparisons correction at each frequency and time-point, yielding similar results.

To further compare EEG power spectral density between pre, during and post-events, for ALA and control stare conditions, we used paired t-tests at each frequency in MATLAB; p-values were corrected across frequencies. Similarly, we compared Φ* values across time, with paired t-tests between the bin with maximum Φ*, and all other time-points; p-values were corrected across time. We evaluated machine learning accuracy using a one-sample t-test, comparing results to chance level performance (50%) at each sample across time; p-values were corrected across time.

We compared other effects using linear models (LM in R), reporting t statistics and corrected p-values for all relevant main and interaction effects, or correlations (cor in R), reporting correlation coefficients and p-values. For behavioral changes, we compared data across condition (During versus Pre/Post) for ALA events. For Φ* controls (Fig. 5*C,D*), we compared data between task condition (resting wakefulness versus fixation task) for all non-stimulation samples. Statistical analysis was then performed separately for each stimulation condition (10/50/200Hz stim-on compared to stim-off). We normalized spike rates to the pre-event condition, then tested with linear models to produce an intercept and p-value. P-values were corrected for all brain regions in the same condition.

For *H* and Φ*, we centered values relative to those obtained during fixation controls to account for day-to-day variations. Condition (*C*, During vs Pre/Post) and event (*E*, ALA vs control stare) were coded as centered, dichotomous variables. We included animal (*A*) as a covariate, though similar results were obtained without its inclusion.

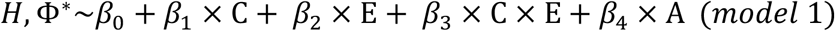

For long-duration stimulation effects, we tested the relationship between local event probability (*LEP*) and local stimulation predominance (*LSP*), first using the cor function in R, followed by linear modeling to produce best-fit lines. Both variables were normalized to their maximum, thus ranging between 0 and 1. Comparisons between ALA and control stare conditions used the following interaction model, where event (*E*) was a centered, dichotomous variable (ALA versus control stare). Animal (*A*) was included as a covariate, though results were similar either way.

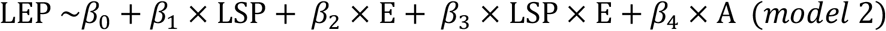

This effect could be simplified by comparing the event probability according to stimulation series type (*SS*, Stim versus No-Stim). Event (*E*) was a centered, dichotomous variable (ALA versus control stare); and animal (*A*) was included as a covariate, though results were similar either way.

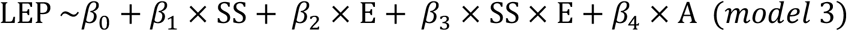

To account for differences seen in the transition between stimulation and non-stimulation series, relative to the established phase, we updated the model to include a 3-way interaction between *SS, E*, and establishment (*EST*, Transition versus Established), as a centered, dichotomous variable. Simple effects were consistent between the two models.

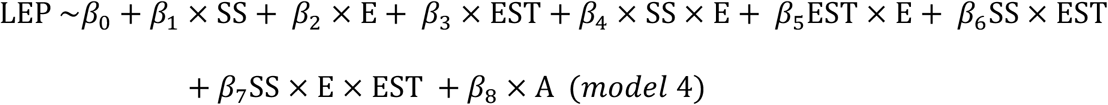

We compared MDA results along the dimensions of frequency and brain area using a dummy coding approach. Patterns were compared between condition (*C*, during versus pre) with the following models. P-values were corrected across all frequency and area comparisons.

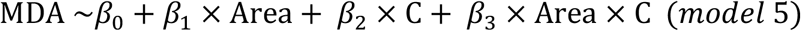

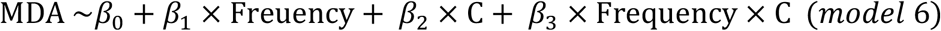

## RESULTS

### Thalamic stimulation during wakefulness reduces behavioral and EEG signatures of consciousness

We simultaneously stimulated across the dorsal-ventral extent of CL in awake monkeys via 16 contacts of a linear electrode array (Fig. 1). On a subset of recording blocks (17.75% of all non-fixation task blocks, including those with and without stimulation; stimulation frequency-and history-dependence detailed below), animals went from normal behaviors, including scanning the room and making movements with their face and limbs, to suddenly holding their gaze and staring vacantly (Fig. 4*A*). These events occurred without any overt sensory cause, such as a noise that might draw the animal’s attention, or visual cue. Animals became still and would hold their gaze for an abnormally long period of time (> 4.2s, M = 17.64s, SD = 5.96s, 99.93 percentile of all stable fixations which lasted at least 1s). Electrophysiological examination of these events also revealed transient increases in lower frequency (1-9Hz) activity, especially in the EEG. This syndrome resembled absence epilepsy; we even observed occasional subtle eye flutters and small lip movements similar to those often seen in clinical patients during the events (Blumenfeld, 2005; Hughes, 2009; Tenney and Glauser, 2013). Given absence seizures have been reported to result from thalamic DBS in cats (Jasper and Droogleever-Fortuyn, 1948; Hunter and Jasper, 1949; Ingvar, 1955) and monkeys (Steriade, 1974), we termed this phenomenon absence-like activity (ALA).

ALA events (N = 60) corresponded to substantial reductions in eye movement (Fig. 4*B*, t(55) = 9.05, n = 56, p = 1.75*10-12, paired t-test), mouth movements (Fig. 4*D*, t(56) = 9.05, n = 57, p = 4.95*10^−7^, paired t-test), and nose movements (Fig. 4*E*, t(56) = 4.99, n = 57, p = 6.18*10^−6^, paired t-test) relative to pre and post ALA conditions. Eye lids remained open with only small amplitude changes during ALA compared to pre/post epochs (Fig. 4*C*, t(56) = 13.72, n = 57, p = 2.00*10^−16^, paired t-test). During ALA, the EEG showed substantial increases in low frequency power relative to the pre (Fig. 4*F* top, 3-22 Hz, |t(1798)| ≥3.43, n = 1800, p_c_ ≤ .0465, paired t-tests) and post condition (Fig. 4*F* top, 6-22 Hz, |t(1798)| ≥4.47, n = 1800, p_c_ ≤ .0004, paired t-tests). EEG power prior to and after ALA are similar to each other (Fig. 4*F* top), except for small differences in the δ band (1-4 Hz, |t(1798)| ≥3.69, n = 1800, p_c_ ≤ .0204, paired t-tests). These results cannot be explained by decreased behavioral activity alone. Control stares (STR), behaviorally similar long-duration fixations (N = 64, M = 15.31s, SD = 6.63s, only extreme examples were selected to best match ALA durations) were similar in power relative to pre and post conditions (Fig. 4*F* bottom), except that STR events tended to have lower delta power relative to pre (0-5 Hz, |t(1918)| ≥4.05, n = 1920, p_c_ ≤ .0046, paired t-tests) and post (0-7 Hz, |t(1918)| ≥4.72, n = 1920, p_c_ ≤ .0004, paired t-tests) conditions. Substantial changes are not present in the EEG between ALA and control stares until around 1.5 seconds after the behavioral onset of an event (Fig. 4*G*), and include significant clusters of increased power only at lower frequencies (δ, θ and α bands, 0-15 Hz, Tsum = 948.889, p = .0130, Nboot = 10,000, permutation test).

### ALA corresponds to selective reductions in neural complexity and integration

Measures of complexity are promising neural indicators of consciousness for both animal models and human subjects (Seth et al., 2008; Haun et al., 2017; Arsiwalla and Verschure, 2018; Afrasiabi et al., 2021). We computed entropy (*H)*, a measure of the complexity of neural interactions, and Φ*, which additionally measures integration of network parts, i.e., how partitioning the network influences the generation of information. We compared how *H* and F^*^ associated with consciousness before, during, and after ALA events and control stares, as well as during resting wake epochs, the fixation task, and two types of general anesthesia (Isoflurane and propofol). We have previously shown that both F^*^ and *H* tend to be significantly reduced during anesthesia or sleep compared to wakefulness (Afrasiabi et al., 2021). Relative to the levels recorded during the fixation task, both *H* (Fig. 5*A*) and Φ* (Fig. 5*B*) were decreased during ALA, but not control stares, compared to pre and post conditions. The interaction of condition (During vs Pre/Post) and event (ALA vs control stare) was significant for both *H* (F(295) = 14.578, n = 300, p = 1.64×10^−4^, ANCOVA) and F^*^ (F(295) = 34.261, n = 300, p = 1.28×10^−8^, ANCOVA). Differences in Φ* cannot be explained by general effects of thalamic DBS. When excluding ALA events, there were no significant differences in Φ* between stim-on (S+) or stim-off (S-) epochs (Fig. 5*C,D*) at any stimulation frequency (t_50Hz_(62) = .29, n = 64, p = .773; t_10Hz_(88) = .65, n = 90, p = .517; t_200Hz_(114) = .732, n = 116, p = .466, paired t-tests). Φ* also proved statistically similar between resting wake and on-task fixation conditions (Fig. 5*C,D*, t(118) = 0.925, n = 120, p = .357, paired t-test), meaning decreases in Φ* were selective to ALA. These results support the conclusion that decreases in Φ* during ALA events indicate an event-specific reduction in consciousness.

### Changes in consciousness correspond to changes in cortico-striatal associations

Calculating Φ* requires computing the minimum information partition (MIP), the cut in the system resulting in the minimum amount of information loss. This indicates system parts which are most integrated. We compared the distribution of MIPs for the largest consistent subsystem (containing CN, layers of frontal and/or parietal cortex) across different states of consciousness (isoflurane/propofol anesthesia, resting wake, fixation, pre/post/during ALA and control stares) using kernel density estimates and two-sample Kolmogorov–Smirnov tests (Fig. 5*E,F,G*). The results revealed that ALA events exhibit a substantially different MIP distribution relative to pre (KS(28) = 0.80, n = 30, p_c_ = 1.10×10^−10^, KS-test) and post (KS(28) = 0.50, n = 30, p_c_ = 0.0012, KS-test) conditions, while control stare events did not. ALA distributions were clearly different from control stares (KS(28) = 0.65, n = 30, p_c_ = 9.63×10^−7^, KS-test), as well as resting wakefulness (KS(28) = 0.85, n = 30, p_c_ = 3.70×10^−12^, KS-test), and the fixation task (KS(28) = 0.83, n = 30, p_c_ = 2.06×10^−11^, KS-test). But ALA distributions were similar to distributions under isoflurane (KS(28) = 0.3, n = 30, p_c_ = .8195, KS-test) and propofol (KS(28) = 0.18, n = 30, p_c_ = 1, KS-test) anesthesia. See Figure 5G for full KS-test comparisons. This shows that the network is reconfigured in the same way for ALA as for general anesthesia.

To further interrogate this effect, we used the MIP distributions to determine the probability of different areas associating on the same side of the MIP as a measure of connection strength (Fig. 6*A,B*). While cortical connections remained relatively consistent across conditions, CN commonly associated with parietal regions in pre/post-ALA, STR, resting wakefulness and fixation task (Fig. 6*A*, red dashed border), but associated with frontal regions in ALA, isoflurane and propofol (Fig. 6*B*, blue dashed border). We have previously shown a similar switch in configuration to reflect changes in consciousness (Afrasiabi et al., 2021). These configuration differences were substantial relative to those expected by chance. Ld-CN connection differences were especially strong and significant (|Z| ≥ 2.15, p_c_ ≤ .0312, Nboot = 10000, permutation test) compared to a null distribution (Fig. 6*C*). To further link this effect to consciousness, we simulated a null distribution representing consciousness by randomly selecting combinations of MIPs found in the wake or fixation task conditions. For ALA, isoflurane and propofol, the associations of CN with cortical areas Ld, Ls, Fd, and Fs were all substantially different (|Z| ≥ 6.62, p ≤ 3.62×10^−11^, Nboot = 10000, permutation test) relative to the expected distribution under consciousness (Fig. 6*D*). These results imply that network-level activity during ALA events shifts into a configuration indicative of a reduction in consciousness.

We computed Φ* in 1s sliding windows (.1s steps) leading up to, and beyond, the behavioral onset of ALA and control stare events (Fig. 6 *E,F*). We analyzed data 10.5 seconds prior to event onset, and up to 4.5 seconds after, as these limits represented the maximum consistent duration across all events. On average, Φ* began to decrease about 3.0s before ALA onset (Fig. 6*E*), when the MIP configuration switched from what is commonly seen in conscious states to the MIP most favored under anesthesia (Fig. 6*F*). This change in MIP configuration continued until 2.9s after ALA onset when it switched back to the wake-favoring MIP. Following the MIP switch, Φ* was significantly lower than the maximum value between -1.6 and 3.9s relative to event onset (|t(82)| ≥ 3.72, n = 84, p_c_ ≤ 0.049, paired t-tests) for ALA, but was stable during control stares and never significantly lower than the maximum (Fig. 6*E*). Similarly, the MIP configuration during control stares remained in the wake-favoring configuration throughout and never changed relative to event onset (Fig. 6*F*). These results imply that the decreases in consciousness associated with ALA predate the onset of abnormal behavior by 1-3s.

### ALA, not control stares, are modulated by thalamic DBS in a frequency-dependent manner across acute and long-term time scales

We argue below that thalamic DBS (specifically at 10 and 200Hz) induces atypical network dynamics biasing towards ALA. That is, not all ALA events occurred as the direct result of an individual DBS stimulation, but rather, ALA in general is modulated by the entire experimental history of DBS in a frequency-dependent manner. We performed 338 experiment blocks in the resting wake condition. ALA events were significantly more common during recording blocks that included thalamic DBS (23.13%, n = 37) than those without (12.92%, n = 23, Z = -2.45, p = .014). Each block involved periods of no stimulation with three replicated stimulation epochs (Fig. 2). ALA was significantly more likely to occur within the bounds of the stimulations (64.86%, n = 24) than outside (35.14%, n = 13) compared to a null distribution representing random occurrence (Z = 2.36, p_c_ = .036, n = 37, permutation test). Moreover, ALA events were overrepresented near the onset (Fig. 7*A*) of a stimulation epoch (within 2s of stimulation onset, 25%, n = 6; Z = -2.58, p = .031, Nboot = 100,000, permutation test). These results imply that thalamic DBS serves as a trigger for ALA under certain conditions.

In line with our hypotheses, ALA probability was higher with 10Hz (23.9%) and 200Hz (30.0%) stimulations, than it was for 50Hz (14.8%) and no-stimulation controls (12.9%) (Fig. 7*B*). Permutation tests revealed that 200Hz stimulations gave rise to significantly increased probability of ALA (Z = 2.71, p_c_ = .025, Nboot = 100,000, permutation test) compared to the average occurrence (17.8%), while experiments without stimulation (NS) had a significantly decreased probability (Z = -2.44, p_c_ = .044, Nboot = 100,000, permutation test). We found similar results with causal power (CP, Fig. 7*D-F*), a probability ratio indicating the strength of causal relationship between two variables relative to alternatives (see METHODS). Positive or negative CP indicates that a given cause (stimulation condition) increases or decreases effect likelihood (ALA occurrence) more so than other known plausible causes (Griffiths and Tenenbaum, 2005). Across the experimental history for both monkeys, CP was typically positive for 200Hz and 10Hz stimulations, and negative for no stimulation and 50Hz stimulation (Fig. 7*C,D*). Simulated null-distribution tests indicated causal power for ALA was significantly positive for 200 Hz stimulations (ΔCP = .175, Z = 2.66, c = .031, Nboot = 100,000, permutation test), and significantly negative for no stimulation (ΔCP = -.133, Z = -2.58, p = .031, Nboot = 100,000, permutation test) relative to expectations under a null distribution. These results imply that abnormally high frequency activity applied to CL promotes ALA with our DBS method.

The occurrence of ALA also appeared to depend on the stimulation order. In our study, we pseudorandomly assigned the order of stimulation experiments between no-stimulation control, 10Hz, 50Hz and 200Hz conditions (Fig. 2) within each experimental run. Thus, for established days in a stimulation series, we could compare the probability of ALA occurring during blocks with a given stimulation frequency relative to that of the preceding stimulation (Fig. 7*F*). Interestingly, 50Hz stimulations, while the least likely to produce ALA, were more likely to do so if preceded by a 200Hz (CP = .091) rather than 10Hz stimulation (CP = -.147). 10Hz stimulations more commonly exhibited ALA after 200Hz (CP = .063) relative to 50Hz (CP = -.273) stimulations; and 200Hz stimulations more commonly exhibited ALA after 10Hz (CP = .515) relative to 50Hz (CP = .038) stimulations. Although none of these effects reached significance with permutation tests (due to the reduced sample sizes when considering all possible combinations of stimulation conditions), the synergistic effect of 10Hz on 200Hz was trending (Z = 1.63, p = .102, Nboot = 100,000, permutation test). While the limited power of these analysis does not warrant strong claims, it is still interesting to note that abnormal stimulation frequencies relative to CL (10Hz and 200Hz) both tend to exacerbate ALA and have positive CP. In comparison, 50 Hz stimulations, which we have previously shown to increase consciousness in macaques (Redinbaugh et al., 2020), tend to have negative CP and decrease ALA occurrence.

Irrespective of stimulation frequency, ALA probability increased with the proportion of DBS blocks over consecutive days (Fig. 7*G*). When computing the probability of ALA in sliding windows across 32 blocks (approximating 4 experimental runs and 2 recording days), the local probability of ALA was positively correlated (Fig. 7*G*) with the predominance of stimulations within the same window (r = .5105, t(309) = 10.40, p = 6.54×10^−16^). Control stares were not significantly correlated with stimulations (r = -.0466, t(309) = -.82, p = .4141). An interaction model verified that the effect of stimulation predominance on event probability was substantially different between ALA and control stares (F(613) = 68.043, n = 618, p = 9.79×10^−16^, ANCOVA). Our experimental paradigm consisted of shifting between series of recording days that either included stimulation or did not, and each series typically lasted for at least one week, with the first two days of a series being considered transitional (Fig. 2). ALA on no-stimulation experimental runs was less common (38.33%, n = 23) than those with stimulation (61.67%, n = 37; Z = -2.45, p = .014, Nboot = 100,000, permutation test). Further, local event probability was significantly higher for experimental runs that included thalamic DBS (Fig. 7*H*, “All” bars) compared to those that did not (t(613) = 6.639, p = 6.96×10^−11^, n = 618, t-test). This effect was substantially larger than the same effect computed for stare controls, yielding a significant interaction (F(618) = 18.787, n = 618, p = 1.71×10^−5^, ANCOVA). Interestingly, local ALA probability was higher in the transition period for no-stimulation series, i.e., when the previous few recording days had included thalamic DBS, and decreased markedly after the series was established (Fig. 7*H*). In contrast, local ALA probability was reduced when transitioning from a no-stimulation into a stimulation series, and increased after the series was established. A 3-way ANCOVA verified these effects were significant and specific to ALA events rather than stare controls; the three-way interaction of experimental series type (Stim, No-Stim), establishment (Transition, Established), and event (ALA vs Control Stare) was significant (F(609) = 92.294, n = 618, p = 2.00×10^−16^, ANCOVA). These results imply that thalamic DBS influences the prevalence of ALA over longer periods of time. Thus, while DBS can directly initiate ALA events, it can also produce longer-duration vulnerabilities that gradually dissipate after stimulation experiments end.

We also included a fixation task in this study as a control (331 experimental blocks recorded), where DBS did not have the same effect. ALA only occurred on 2.72% of fixation task blocks compared to 17.75% of resting wake blocks; this difference was significant (Z = 6.389, p = 1.67×10^−10^, Nboot = 100,000, permutation test). Interestingly, this finding matches the symptomatology of absence epilepsy, as seizures tend to decrease with cognitive engagement in animal models (Van Luijtelaar et al., 1991).

### ALA events correspond to reduced neural firing

We compared the firing rate of the same neurons, normalized to the pre-condition, across ALA, control stare, post, resting wake and fixation task epochs (although this reduced the relatively small sample size further, it better enabled comparisons with all controls). Firing rates were very similar to the pre-condition for all conditions except ALA (Fig. 8). While all areas had lower firing on average during ALA, this difference was only significant for Fs (t(22) = -4.16, n = 23, p_c_ = .002), Ld (t(32) = -3.06, n = 33, p_c_ = .022), and CN (t(13) = -3.22, n = 14, p_c_ = .027). The effect was trending in CL (with a relatively small sample size, as CL recordings only occurred during experiments without thalamic DBS) after multiple comparisons control (t(25) = -2.26, n = 26, p = .033, p_c_ = .098). These results could reflect changes in neural gain/excitability, or reduced drive along specific input pathways.

**Figure 8.**
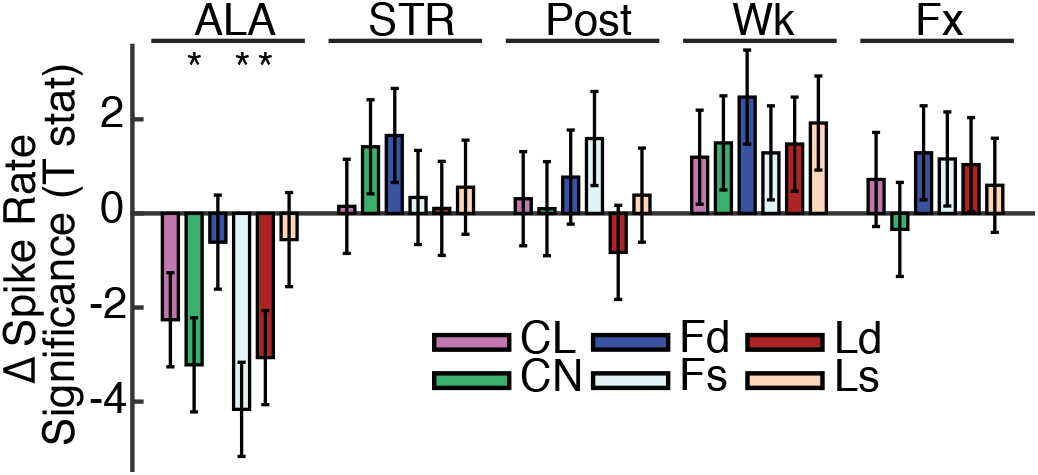
Neural firing rates decrease during ALA relative to control conditions. Spike rate differences normalized to the pre-condition (± SE of the point estimate) for different brain areas under ALA, control stare (STR), post, resting wake (RW), and fixation (FX) conditions. Neurons were analyzed in resting wake, fixation, and pre/post-event epochs only for neurons recorded during ALA or STR events (N_CL_ = 60, N_CN_ = 19, N_Fd_ = 15, N_Fs_ = 38, N_Ld_ = 46, N_Ls_ = 21).

### Reduced consciousness in ALA is associated with increases in low-frequency power spectral density

To further evaluate neural signatures of decreased consciousness, we compared LFP spectrograms in different brain regions between ALA and control stare conditions relative to event onset. In general, power tended to be lower 3-5s prior to ALA onset (Fig. 9*A-F*). This was strongest in CL (Fig. 9*F*) which had a significant cluster from θ-*γ*_H_ frequencies from approximately -5 to -1.8s relative to event onset (Tsum = -650.76, p = 9.99×10^−5^, Nboot = 10,000). Power tended to increase in most areas at low frequencies just prior to event onset, and remain elevated at lower frequencies, especially in Ls (Fig. 9*B*) and Ld (Fig. 9*D*), as the event continued. Power in CL tended to be much higher -1 to 4s around onset, especially at θ and α frequencies. Figure 9*F* shows the significant cluster (Tsum = 282.74, p = 0.0013, Nboot = 10,000). See Table 1 for full statistical results.

**Table 1.** Statistical results of cluster analysis for LFP power spectral density time-frequency spectrograms. Area identity (TP), N (ALA, control stare), cluster identity, T_sum_ statistic, and p-value for each significant cluster as identified in Figure 9. Positive T_sum_ indicates ALA power spectral density > control stare.

| Tp | N | Clust | $T_{sum}$ | p | Tp | N | Clust | $T_{sum}$ | p | Tp | N | Clust | $T_{sum}$ | p |
| --- | --- | --- | --- | --- | --- | --- | --- | --- | --- | --- | --- | --- | --- | --- |
| CL | 345, 506 | 1 | -650.76 | 0.0001 | FEFd | 331,291 | 1 | -61.94 | 0.0328 | FEFs | 451, 445 | 1 | -60.18 | 0.0296 |
|  |  | 2 | -122.71 | 0.0347 |  |  | 2 | -52.51 | 0.0494 |  |  | 2 | -107.86 | 0.0035 |
|  |  | 3 | 282.74 | 0.0013 |  |  | 3 | -60.74 | 0.0343 | LIPs | 492, 525 | 1 | 197.50 | 0.0077 |
|  |  | 4 | -143.94 | 0.0222 | LIPd | 572, 548 | 1 | 183.84 | 0.0063 | CN | 1156, 1149 | 1 | -81.64 | 0.0003 |

**Figure 9.**
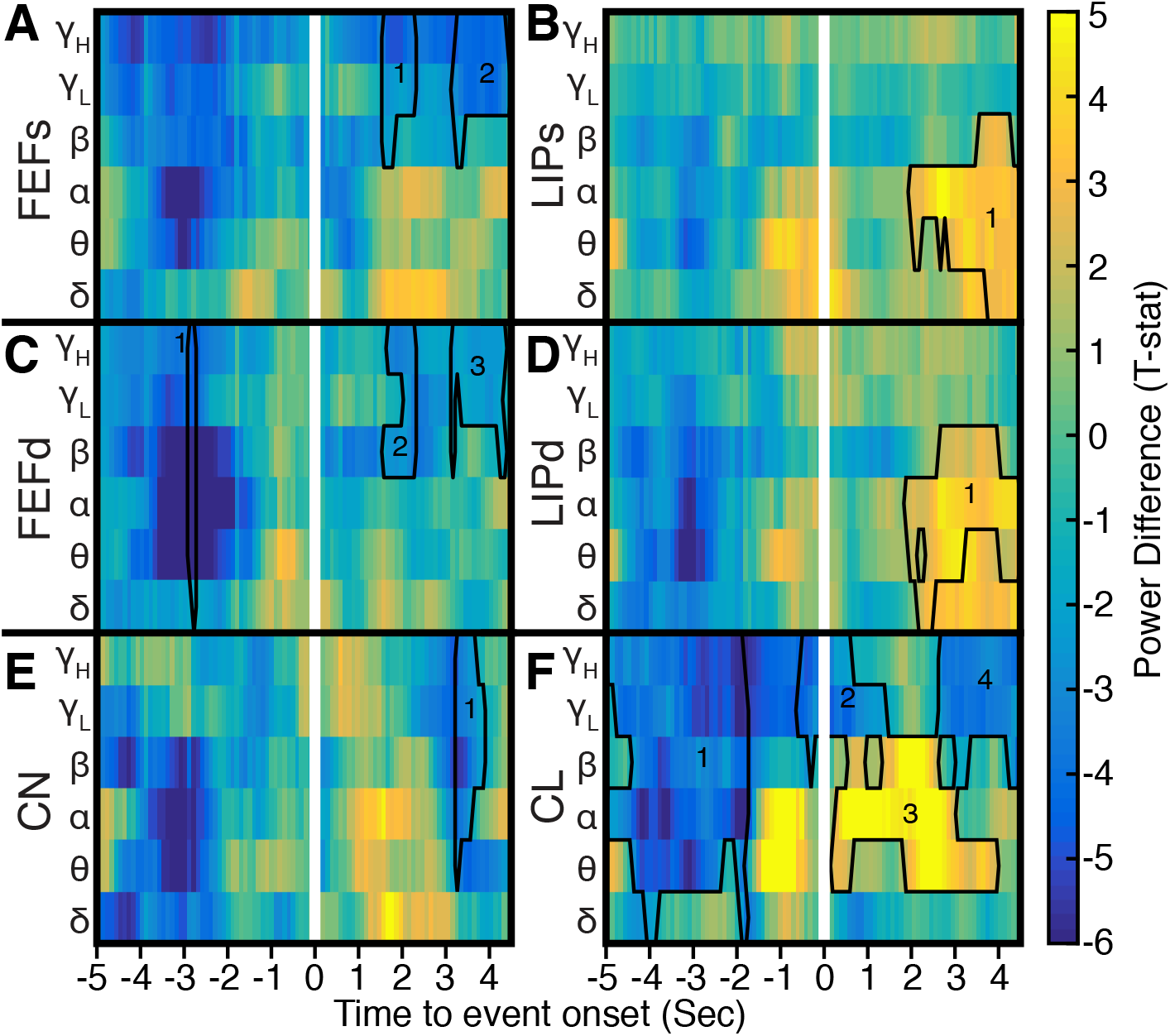
ALA is associated with increases in low-frequency power relative to control stares. **A-F** Spectrograms of power differences (ALA – control stare) for each frequency band across time aligned to event onset (white vertical line). Results include all recorded ALA and control stares for (**A**) superficial FEF, (**B**) superficial LIP, (**C**) deep FEF, (**D**) deep LIP, (**E**) CN, and (**F**) only the subset of events without thalamic DBS for CL. Significant clusters are outlined in black with numerical labels referenced in Table 1 for statistical details.

To further investigate the changes in power spectral density influencing ALA, we trained a Bayesian classifier to differentiate between ALA events and control stares, using the LFP power in different frequency bands (δ, θ, α, β, and *γ*_L_) from CN, Ld, Ls, Fd, and Fs calculated in 1s sliding windows (0.5s stride) relative to event onset. CL was excluded because it was only available on a subset of samples, hindering convergence of the parameter estimators. Decoding was significant (Fig. 10*A*) as early as 4s prior to event onset and remained significant thereafter (|t_-4 to4.5_(9)|≥4.66, n = 10, pc≤.0272). This window was very similar to the time course of changes in Φ* (Fig. 6*E,F*). We used mean decrease in accuracy (MDA) analysis to estimate which of the LFP features (5 areas x 5 frequency bands) contributed most to decoding accuracy across time (Fig. 10*B*). The MDA is computed by comparing accuracy when the machine learning model includes all possible features to an impoverished model where the information content of a feature has been negated by permuting the sample orders. The greater the MDA, the greater the contribution of that feature. Prior to event onset (−2.5 to -0.5s, Fig. 10*A*, blue horizontal bar), exclusion of CN_β_, Ld_θ,α,β_, or Ls_θ,α_ resulted in the largest decreases in accuracy. Overall, features from Ld (Fig. 10*C*, |t(1998)| ≥ 253, n = 2000, p_c_ ≤ 1.4×10^−15^), and features related to β power (Fig. 10*D*, |t(1998)| ≥ 253, n = 2000, p_c_ ≤ 1.4×10^−15^) conveyed more information than others before events.

**Figure 10.**
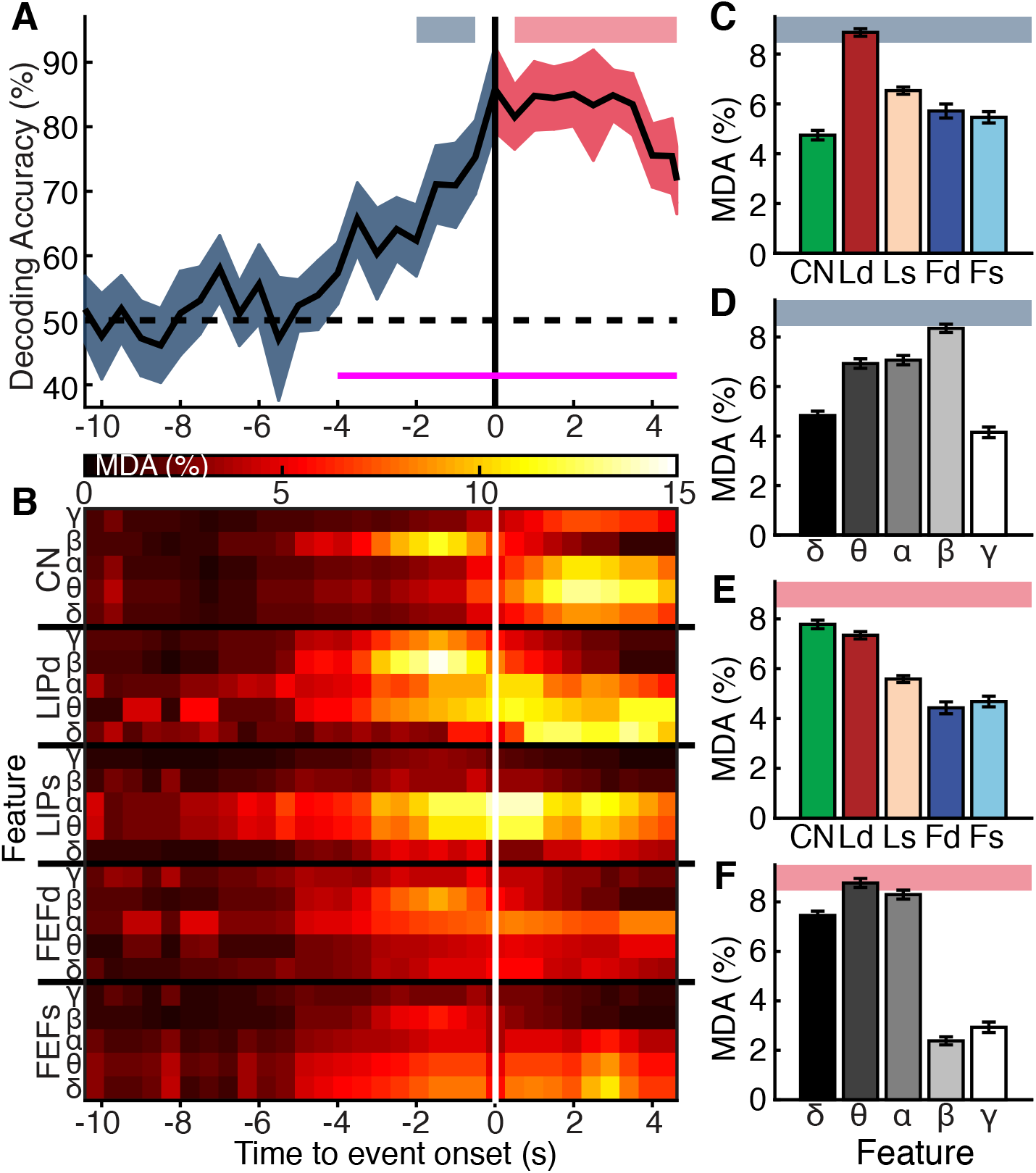
ALA and stare controls are readily decodable on a similar time-frame as changes in consciousness, and represent a clear shift in the relative importance of different frequencies. ***A***, Decoding accuracy (±SE) across time relative to event onset of ALA and stare controls using 25 frequency (δ, θ, α, β, *γ*) by brain area (CN, Ld, Ls, Fd, Fs) features. Thin saturated pink line shows period of significance when decoding accuracy is greater than chance (50%). ***B*** Mean decrease in accuracy (MDA) attributed to each model feature across time. Higher values indicate higher feature importance. **D**-**F**, MDAs further analyzed for the period of time just prior to event onset (thick blue horizontal bar in (**A**)) for features by (**C**) area and (**D)** frequency, and for the period of time just after ALA onset (thick red horizontal bar in (**A**)) for features by (**E**) area and (**F**) frequency. Bar graphs show average MDA (± SD).

Between .5 to 4.5s following ALA onset (Fig. 10*A*, thick red horizontal bar), feature importance shifted to lower frequencies. Feature importance was significantly higher for δ, θ, and α frequencies, and lower for β and *μ* frequencies after the shift (|t(1998)| ≥ 127.3, n = 2000, p_c_ ≤ 1.0×10^−15^). The importance of CN significantly increased, and importance decreased for all other areas (|t(1998)| ≥ 78.75, n = 2000, p_c_ ≤ 1.0×10^−15^). Overall, features from CN and Ld (Fig. 10*E*, |t(1998)| ≥ 210.56, n = 2000, p_c_ ≤ 2.0×10^−15^), and features related to δ, θ, and α power (Fig. 10*F*, |t(1998)| ≥ 552.2, n = 2000, p_c_ ≤ 1.6×10^−15^) conveyed more information than others (see Fig. 10*B* for the most important individual features, i.e., CN_d,θ,α_, Ld_d,θ_, Ls_θ,α_, and Fs_d_). These results indicate that ALA is associated with a substantial increase in low frequency activity between 1-15 Hz. As this activity is most substantial and informative when found in subcortical and parietal structures, these regions may play the largest role in inducing and promoting ALA.

### Reduced consciousness in ALA is associated with changes in CST functional connectivity

To evaluate functional connectivity signatures associated with reduced consciousness, we compared LFP-LFP coherence between the different brain regions representing anatomically-motivated pathways (Fig. 11*A*) relative to event onset between ALA and STR conditions. Changes in coherence were substantial and dynamic across time, leading to a large number of significant clusters (Table 2) in time-frequency coherograms. Thalamo-cortical coherence (Fig. 11*B,C*) increased transiently at d about 3.5s before onset, followed by θ and α about 2.5s before onset (CL-Fs cluster 3, CL-Fd clusters 4 and 5, CL-Ls cluster 2, and, CL-Ld cluster 4, Tsum ≥ 43.23, p ≤ .0084, Nboot = 10,000). This θ and α coherence only lasted about 1.5s in thalamo-frontal pathways (Fig. 11*B*), but continued in thalamo-parietal pathways (Fig. 11*C*) at least 1.5s after event onset, matching the time course of abnormal θ dynamics exhibited by CL.

**Table 2.**
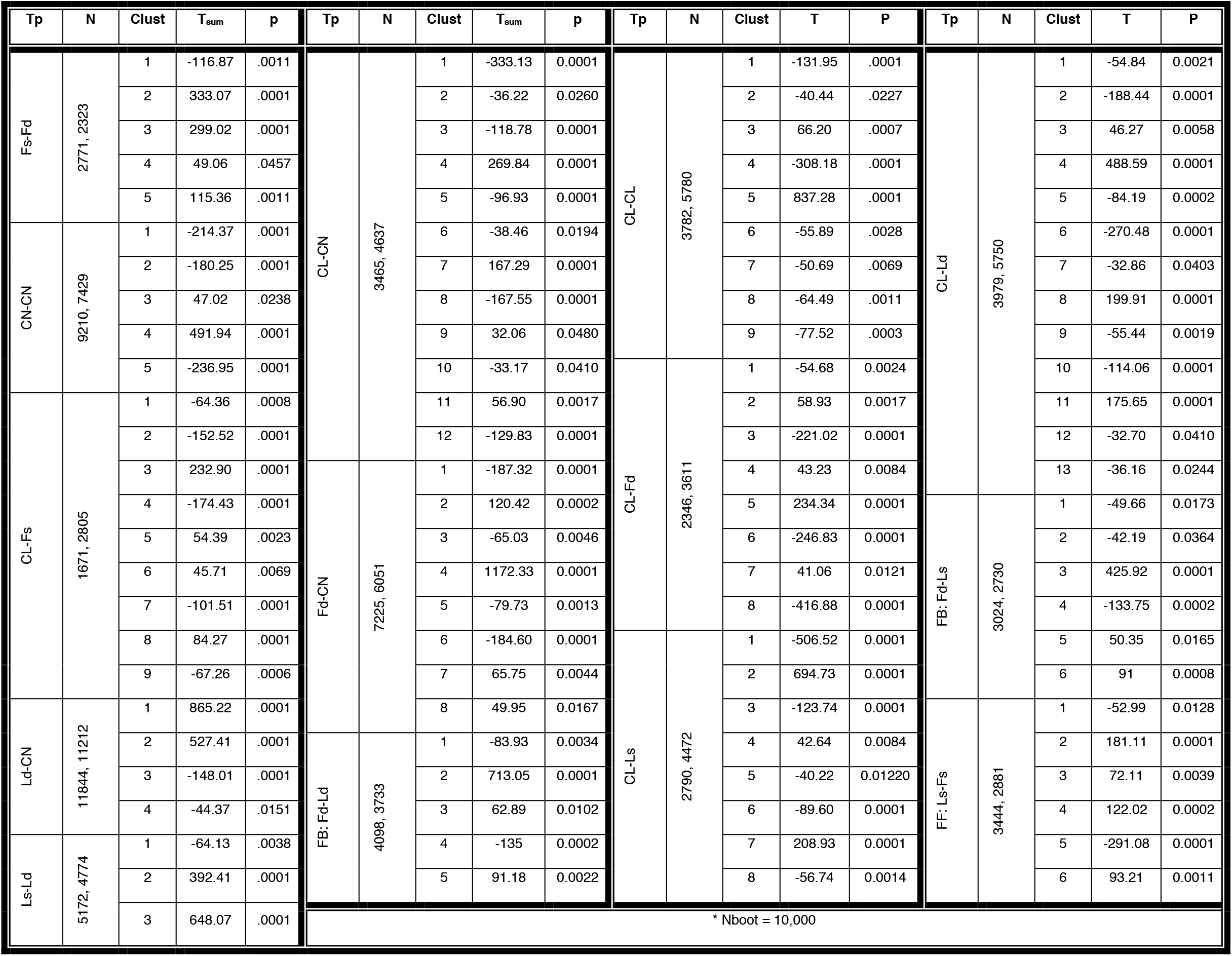
Statistical results of cluster analysis for coherence time-frequency spectrograms. Area identity (Tp), N (ALA, control stare), cluster identity (Clust), T_sum_ statistic, and p-value for each significant cluster as identified in Figure 11. Positive T_sum_ indicates ALA coherence > control stare.

**Figure 11.**
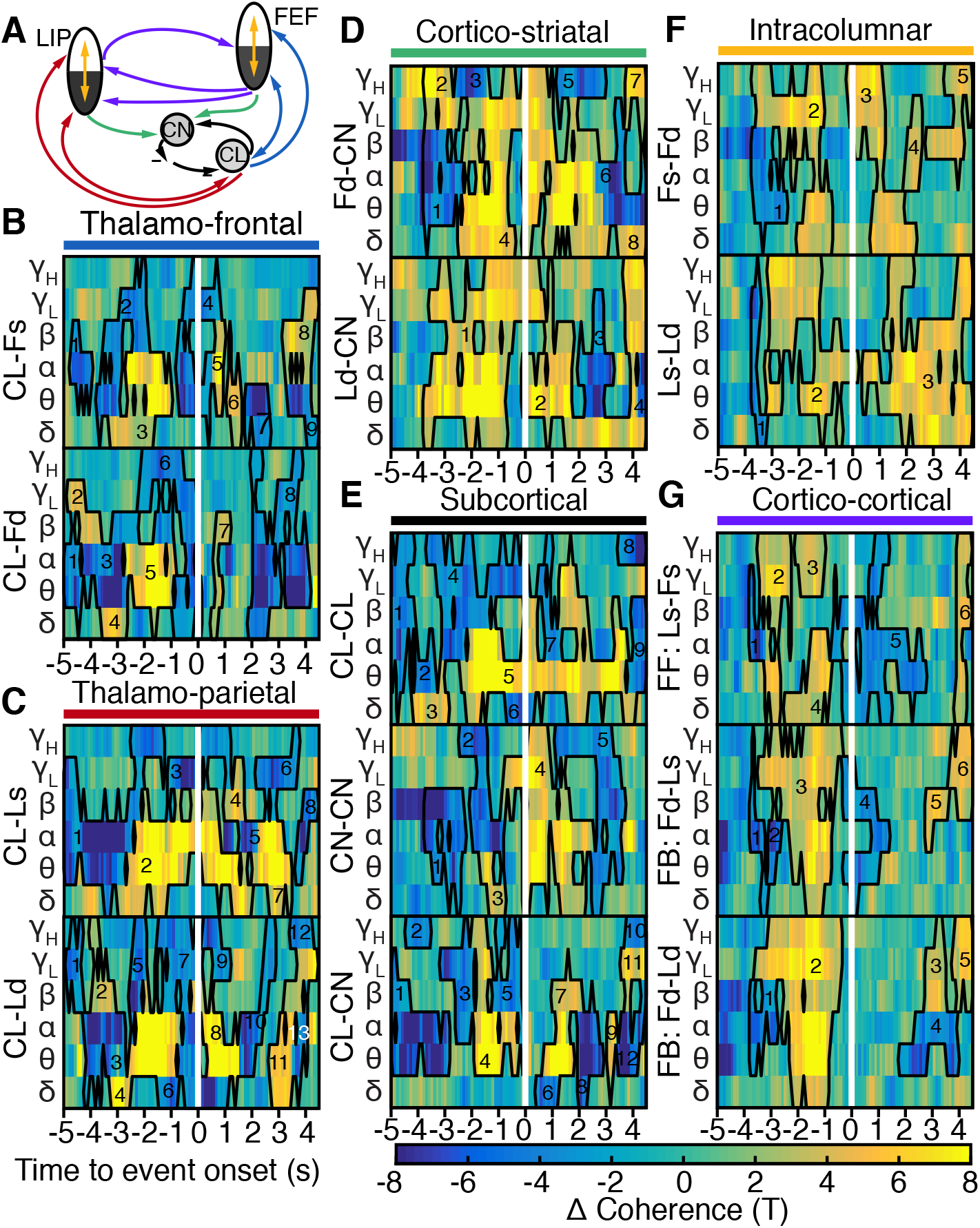
Functional connectivity increases, especially at low frequencies, just before and throughout ALA. ***A***, Diagram showing anatomical pathways isolated in our study. Superficial and deep cortical layers are represented by the respective light and dark regions of the labeled area. Arrow colors relate to the panel subtitle underlines featured in B-G. ***B***-***G***, Time-frequency plots showing coherence differences (ALA – control stare) aligned to event onset (white vertical line) for pairs of LFPs within/between brain area(s). Color scales with the strength of the resulting t-stat. Significant clusters are outlined in black with numerical labels referenced in Table 2 for statistical details. Results are shown for the key (**B**) Thalamo-frontal, (**C**) Thalamo-parietal, (**D**) Cortico-striatal, (**E**) Subcortical, (**F**) Intracolumnar, and (**G**) Cortico-cortical feedforward (FF, Ls-Fs) and feedback (FB, Fd-Ld and Fd-Ls) pathways.

About 3-4s before onset, cortico-striatal coherence (Fig. 11*D*) in θ and α bands decreased along the frontal projection (Fd-CN cluster 1, Tsum = -187.32, p = 0.0001, Nboot = 10,000) and increased along the parietal projection (Ld-CN cluster 1, Tsum = 865.22, p = 0.0001, Nboot = 10,000). Fronto-striatal increases started about a second later, and both fronto-and parieto-striatal coherence persisted, especially at d, throughout the ALA event (Fd-CN cluster 4 and 8, Ld-CN cluster 2, Tsum ≥ 49.95, p ≤ 0.0167, Nboot = 10,000).

Just before ALA onset, CL became internally coherent (Fig. 11*E*) at θ frequencies relative to control stares. This effect lasted from approximately -2.5 to 3.5 s relative to onset (CL-CL cluster 5, Tsum = 837.28, p = 0.001, Nboot = 10,000), matching the time course associated with decreases in Φ*. Thalamo-striatal coherence (Fig. 11*E*) increased at low frequencies (θ, α) just before ALA onset (CL-CN cluster 4, Tsum = 269.84, p = 0.0001, Nboot = 10,000) relative to control stares and remained elevated only briefly thereafter (CL-CN cluster 7, Tsum = 167.29, p = 0.0001, Nboot = 10,000). Striato-striatal coherence (Fig. 11*E*), in comparison, exhibited a transient, broadband increase right at ALA onset (CN-CN cluster 4, Tsum = 491.94, p = 0.0001, Nboot = 10,000).

Intracolumnar coherence (Fig. 11*F*) was initially increased at higher frequencies (β, *γ*_L_, and *γ*_H_) (Fs-Fd cluster 2, Ls-Ld cluster 2, Tsum ≥ 333.07, p ≤ 0.0001, Nboot = 10,000). Coherence then shifted to lower frequencies (δ, θ, and α) just before event onset and continued after ALA onset (Fs-Fd cluster 3, Ls-Ld cluster 3, Tsum ≥ 299.02, p ≤ 0.0001, Nboot = 10,000). These effects appeared to be more consistent in parietal regions relative to frontal. We have previously demonstrated similar shifts in intracolumnar coherence to lower frequencies during general anesthesia (Redinbaugh et al., 2020). This could imply that ALA is associated with perturbation of intracolumnar signaling, thereby contributing to sensory disconnection and reductions in consciousness.

Cortico-cortical interactions involve feedforward (FF) pathways, carrying sensory information from superficial layers of parietal cortex to superficial or middle layers of frontal cortex. Feedback (FB) pathways carry predictions and priorities from the deep layers of frontal cortex to the superficial or deep layers of parietal regions (Markov et al., 2014; Mejias et al., 2016). Around 3s before ALA onset, FF and FB coherence (Fig. 11*G*) showed broadband increases (Ls-Fs clusters 2-4, Fd-Ls cluster 3, Fd-Ld cluster 2, Tsum ≥ 72.11 p ≤ 0.0039, Nboot = 10,000) that dissipated just before event onset (−3 to -1 s). At onset, there were transient broadband decreases in putative Fd-Ls feedback (Fd-Ls cluster 4, Tsum = -133.75, p = 0.0002, Nboot = 10,000), but more persistently, a decrease in putative feedforward Ls-Fs coherence that extended to about 2.5s after onset (Ls-Fs cluster 5, Tsum = -291.08, p = 0.0001, Nboot = 10,000). This latter effect could imply a form of sensory disconnect.

To summarize the complex temporal dynamics across all relevant pathways, we compared the percentage of significant time-frequency tiles (hereon referred to as pixels) within different temporal windows, indicating increases or decreases in high (β,*μ*_L_,*μ*_H_) or low (δ,θ,α) frequency coherence (Fig. 12). This can be considered the predominance of different coherence patterns. Window P1 (Fig. 12*A*, -5 to -2.6s) represents the period of time prior to event onset where decoding accuracy (based on power spectral density) is low and differences in consciousness (Φ*) are insignificant. Significant pixels are rare in this period, indicating similarities in coherence between ALA and control events. Most significant pixels that are present reflect decreases in low-frequency coherence between CL and other regions. Window P2 (Fig. 12*B*, -2.5 to -0.6s) represents the time period prior to event onset where decoding accuracy increases, and decreases in Φ* become significant. This corresponds to an increase in the number of significant pixels indicating increases in low-frequency coherence. These changes span most pathways, but especially those involving CN and deep layers of cortical areas, where the effects are more broadband. At the same time, pixels representing the relationship between CL and other areas reflect high-frequency decreases in coherence, whereas cortico-cortical pathways have increased coherence at higher frequencies predominating.

**Figure 12.**
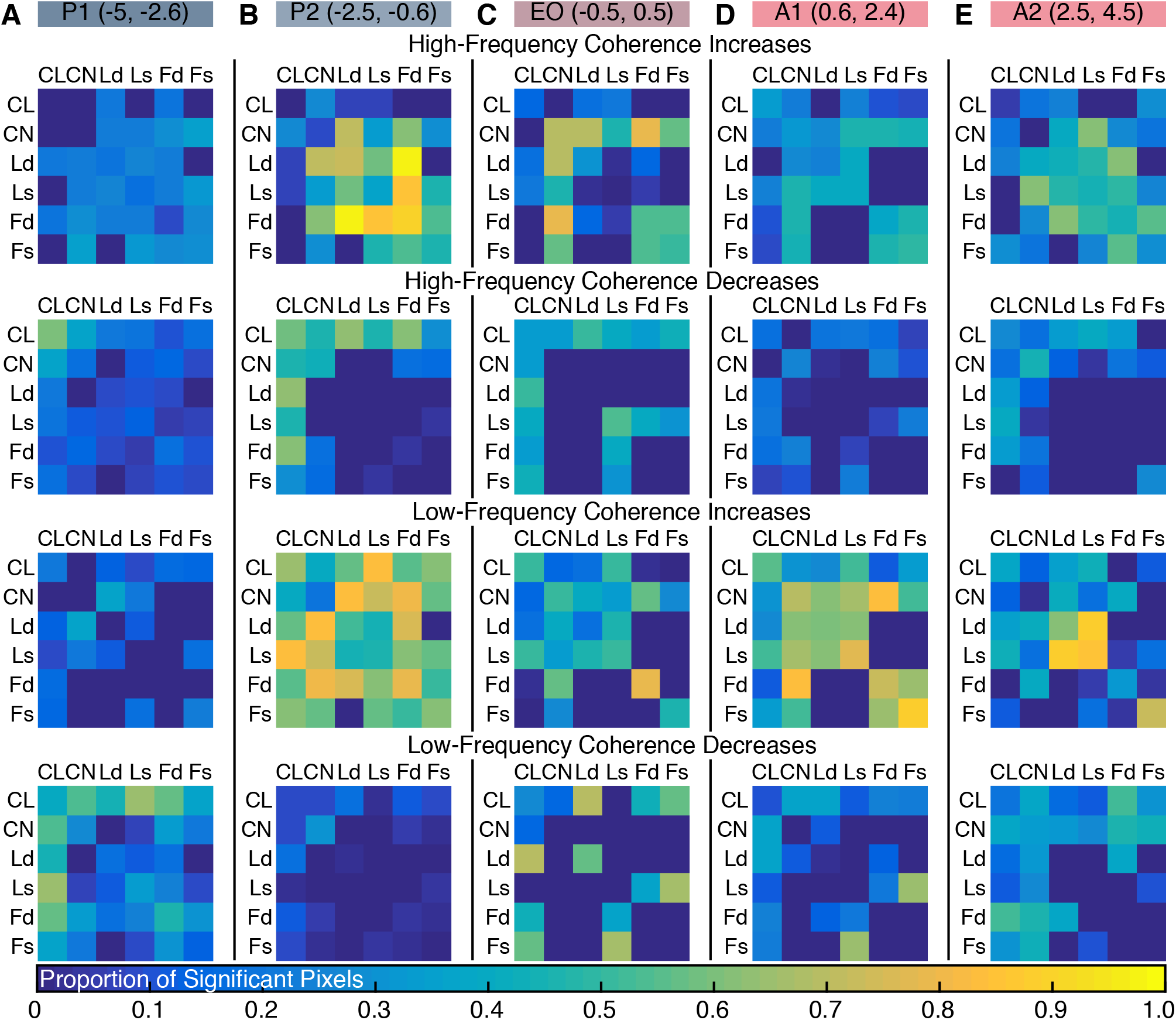
Pathways involving thalamus and striatum dominate the changes in coherence attributed to ALA. ***A***-***E***, For each pairwise comparison across all areas, proportion of significant pixels indicating coherence increases/decreases at higher-frequency (β, *μ*_L_, *μ*_H_) or increases/decreases at lower-frequency (δ, θ, α) in time windows that (**A**) predate significant decoding and changes in consciousness (Pre 1, P1), (**B**) coincide with increased decoding and the start of changes in consciousness (Pre 2, P2), (**C**) coincide with behavioral changes (event onset, EO), (**D**) coincide with sustained changes in behavior and consciousness (After 1, A1), and (**E**) coincide with changed behavior but gradual restoration of consciousness as gauged with Φ* (After 2, A2).

Window EO (Fig. 12*C*, -.5 to .5s) represents the second of data around event onset, where decoding has reached maximum accuracy and changes in Φ* hit the minimum Φ* value. In this window, the majority of pixels representing the relationship between cortex and CN reflect broadband coherence increases. At the same time, there are broadband decreases in coherence predominating for pathways connecting CL to other areas. Window A1 (Fig. 12*D*, .6 to 2.5s) represents the first two seconds after event onset, where decoding remains high and decreases in Φ* remain significant. In this window, there is increased low-frequency coherence within areas (Fig. 12*D*, third row, along diagonal), but not between most areas. Finally, in window A2 (Fig. 12*E*, 2.5 to 4.5s) when behavioral signs of ALA are still present, significant decoding persists and Φ* is gradually returning towards the baseline level, there is predominantly increased low-frequency coherence in parietal cortex, in addition to CL.

## DISCUSSION

We used thalamic DBS to induce acute and spontaneous perturbations in consciousness (ALA events) similar to absence seizures. These events include long-duration, vacant stares coinciding with lower-frequency EEG activity and decreases in neural measures of complexity and integration indicative of consciousness. ALA was best predicted by power changes in parietal cortex and striatum relative to frontal cortex. Just before ALA events, consciousness, as indexed by Φ*, decreased as lower-frequency coherence increased between cortical and subcortical regions (Fig. 13*A*). There was a burst of broadband coherence between the cortex and striatum around ALA onset (Fig. 13*B*). This was followed by dominant lower-frequency coherence within each cortical and subcortical area during ALA (Fig. 13*C*), indicative of increased isolation of individual areas. ALA also involved reduced firing rate in superficial FEF, which could result from reduced input, reflected in the decreased broadband coherence during ALA. We also saw reduced firing rates in LIP deep layers during ALA reminiscent of the reduced excitability of deep-layer cortical neurons previously shown under anesthesia (Redinbaugh et al., 2020; Suzuki and Larkum, 2020), which could result from perturbed intracolumnar communication. Overall, these patterns reflect disruption of typical CST network dynamics, resulting in reduced behavior and consciousness.

**Figure 13.**
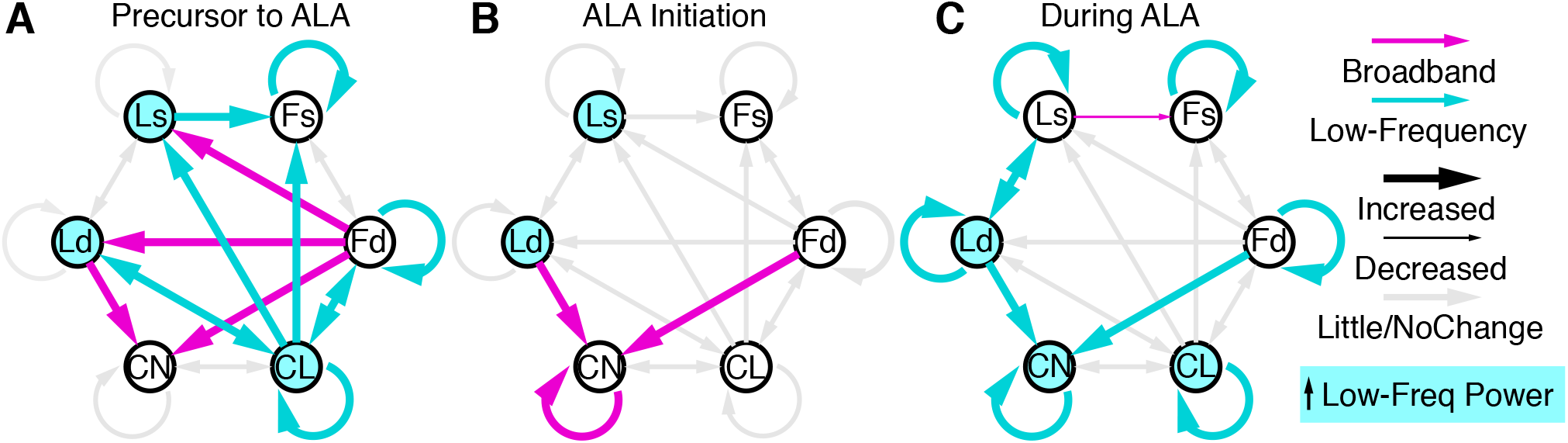
Neural correlates of ALA involve substantial impairment of communication in thalamo-cortical and cortico-striatal circuits. ***A***-***C***, Connection diagrams summarizing key coherence and power patterns within and between different areas (**A**) just before, (**B**) at onset, and (**C**) ongoing during ALA. Arrows show increased (thicker) or decreased (thinner) coherence between specified areas at lower (light blue) or broadband (pink) frequencies. Light gray arrows indicate pathways where coherence changes are relatively small or unchanged. Blue shading in areas shows power increases at low frequencies. Ls and Ld, superficial and deep layers of LIP; Fs and Fd, superficial and deep layers of FEF; CN, caudate nucleus; CL, central lateral thalamus. Diagrams summarize results from Fig. 9-11, and values thresholded at .4 in Fig. 12.

A handful of studies have suggested thalamic DBS can lead to a syndrome similar to absence epilepsy (Jasper and Droogleever-Fortuyn, 1948; Steriade, 1974), and indeed, ALA mirrors the disorder in terms of behavior, decreased consciousness, and network perturbations. Current views attribute absences to disfunction along thalamocortical circuitry (Matricardi et al., 2014). Here, ALA events initiated with increased low-frequency coherence along thalamocortical pathways, especially involving parietal cortex. More recently, absence epilepsy has also been linked to cortico-striatal disfunction (Coenen and Van Luijtelaar, 2003; Berman et al., 2010; Carney et al., 2010; Miyamoto et al., 2019), and rodent models have shown dopaminergic manipulation in the striatum can modulate seizures (Deransart et al., 2000). Again, ALA events initiated with broadband coherence changes along cortico-striatal pathways and involved ongoing decreases in striatal firing rate. Thus, ALA shares key mechanisms with absence epilepsy that could be informative for clinical populations.

The chief distinction between ALA and absence epilepsy is the presence of 3Hz spike-wave patterns in the EEG during absences in humans (Blumenfeld, 2005; Hughes, 2009; Tenney and Glauser, 2013). We observed oscillatory increases during ALA, but at frequencies (3-8Hz) more similar to other reports of absence in macaques (4.5-6Hz) (Marcus and Watson, 1968; Steriade, 1974), or frequencies seen in rodent models (5-10Hz) (Coenen and Van Luijtelaar, 2003). Although we did not detect clear spike-and-wave patterns during ALA, absence seizures can present with abnormal or disorganized morphology more similar to our observations (Sadleir et al., 2006). Further, it has been argued that spike-wave patterns can be found in wild-type mice and may not be related to decreased consciousness associated with absences (Taylor et al., 2019). While this current study does not comment on the consequence of spike-wave patterns, their limited appearance during ALA does imply that symptoms of absence epilepsy can occur by other means.

This study used multi-microelectrode DBS to manipulate consciousness in a frequency-dependent manner. Instead of larger DBS electrodes commonly used clinically, our stimulating electrodes were linear microelectrode arrays designed to conform to CL dimensions in macaques (Fig. 1). Previously, we used this DBS method to increase consciousness in anesthetized macaques in a frequency-dependent manner that was CL-selective and maximally effective at 50Hz (Redinbaugh et al., 2020). This result contrasts with another recent study achieving similar arousing effects using more traditional methodology, but at a much higher stimulation frequency (150Hz) and current (1.0-2.0mA vs our 200*μ*A) (Bastos et al., 2021). These discrepancies could imply a fundamental difference in mechanism between the DBS methods. It is possible that the multi-microelectrode method more directly influences the targeted nucleus in ways that match the stimulation parameters.

This interpretation is supported by our current study, where thalamic DBS modulated ALA at both acute and longer-duration time scales. ALA was most strongly driven by 200Hz or 10Hz perturbations, both of which represent abnormal dynamics for CL in wakefulness, relative to 50Hz stimulation, which produced modulatory effects no stronger than no-stimulation controls. 10Hz is more similar to the dynamics of CL activity in sleep (Steriade et al., 1993; Redinbaugh et al., 2020), while 200Hz represents non-physiological activity or possibly the short inter-spike intervals within bursts featuring in reduced conscious states. In comparison, 50Hz matches the dynamics of fast-firing CL neurons during wakefulness (Glenn and Steriade, 1982; Steriade et al., 1993; Redinbaugh et al., 2020). Clinically, absence seizures are associated with reduced arousal and are more common after sleep deprivation (Sadleir et al., 2011). 10Hz stimulations might reduce arousal and increase vulnerability of the CST system to subsequent 200Hz stimulations. The fact that our results are frequency-specific, and these frequencies are functionally relevant for CL, implies that our method exerts effects by mimicking outputs of CL, and not through antidromic mechanisms.

ALA was associated with stimulation onset, implying that DBS directly influences CST dynamics to promote ALA. Further, ALA became more prevalent as stimulations were performed more frequently, even occurring without DBS application. These results imply that vulnerability to CST circuit disruptions linger and contribute to future disruptions in consciousness. Our stimulations lasted for 1 minute, and were replicated three times per stimulation block, with multiple blocks performed each day. This paradigm could induce neural plasticity (Falowski et al., 2011; Cooperrider et al., 2014) promoting abnormal network dynamics that both exacerbate future responses to DBS and increase the likelihood for resting-state dynamics to spontaneously give way to ALA events. Similar mechanisms seem to contribute to epilepsy, as seizures commonly alter functional connectivity over the course of the disease, with the effect size dependent on seizure rate (Bharath et al., 2015). Interestingly for ALA, plasticity seemed to depend on exposure to the stimulation paradigm, as after experiments ceased, no abnormal events have been observed by experimenters or care staff during everyday behavior.

These results suggest that experimental parameters could be optimized to increase the rate of ALA beyond those we observed. Bilateral stimulation, as used by others (Schiff et al., 2007; Baker et al., 2016), may produce stronger effects than our unilateral approach. Further, our experiments incorporated factors which decreased ALA likelihood (fixation task, no-stimulation blocks, 50Hz stimulations). While these were important controls, future work could optimize effects with repeated batteries of 10Hz and 200Hz stimulations, which trended towards a synergistic effect. Such optimizations increasing ALA likelihood could further enable perceptual experiments in which sensory stimuli are presented during ALA or without perturbing consciousness. Increasing the ALA rate from DBS is likely to also increase the spontaneous ALA rate during non-stimulation transitions, which would further reduce potential confounds, such as artifacts caused by DBS.

This study builds on recent findings about CST circuitry during different conscious states. We previously showed that Φ*, a measure of neural integration, differentiates wakefulness, sleep, and anesthesia (Afrasiabi et al., 2021). Here, Φ* further differentiates resting wakefulness, fixation tasks and stare controls from ALA. The time course of changes in Φ* map onto the time course of decodability of ALA versus control stare events based on cortical and subcortical LFP power, as well as dynamic changes in CST network coherence. In this study, atypical broadband increases in coherence between cortex and striatum were linked to decreased consciousness at ALA onset, just as ongoing low-frequency intra-area coherence was linked to reduced consciousness during ALA events. Critically, this implies that a number of different patterns of aberrant activity can contribute to reduced Φ* and consciousness in CST circuits.

Decreased consciousness also temporally aligned with changes in the integrated structure of the CST network (Fig. 6), such that frontal regions, rather than parietal, became more affiliated with subcortical structures. This finding during ALA generalizes network configuration changes for reduced consciousness under two separate anesthetic agents and, as previously shown, during non-rapid-eye-movement sleep (Afrasiabi et al., 2021). This further challenges cortico-centric views of consciousness (Lamme, 2006; Friston, 2010; Dehaene and Changeux, 2011; Oizumi et al., 2014). Affiliation of parietal and subcortical structures during wakefulness may represent parieto-striatal-thalamic interactions promoting sensory processing and perceptual representations. Shifts of subcortical affiliations to frontal cortex could represent breakdowns in the aforesaid more posterior associative networks and sensory pathways, resulting in decreases in consciousness.

The question remains how disruption of thalamic activity with DBS spreads across the CST network to perturb consciousness. The thalamus can disrupt cortico-striatal interactions by either directly influencing cortex, and consequently cortico-striatal projections, or via thalamo-striatal projections which can directly modulate striatal responses (Kimura et al., 2004). Our results suggest a relatively greater influence of the former. Thalamic DBS caused interference in thalamic outputs which translated to cortico-striatal, more so than thalamo-striatal, perturbances (Fig. 11). Abnormal cortico-striatal inputs would alter basal ganglia output to the thalamus, further perturbing thalamo-cortical, cortico-cortical and cortico-striatal processes. Overall, our results highlight the influential role of the basal ganglia and higher-order thalamic structures, which control information processing and flow through CST networks in ways that support and manipulate consciousness (Schiff, 2010; Fridman et al., 2014).

## ACKNOWLEDGMENTS

Funding: work supported by NIH grants R01MH110311 & P51OD011106, NSF 2021011, BSF grant 201732 and WNPRC pilot grant. We thank D. Cleveland for useful discussions.

